# Metastasis-initiating osteosarcoma subpopulations establish paracrine interactions with both lung and tumor cells to create a metastatic niche

**DOI:** 10.1101/2024.06.09.597967

**Authors:** James B. Reinecke, Amanda Saraf, John Hinckley, Amy C. Gross, Helene Le Pommellette, Leyre Jimenez Garcia, Maren Cam, Matthew V. Cannon, Sophia Vatelle, Berkley E. Gryder, Ruben Dries, Ryan D. Roberts

**Author notes:** **Corresponding author contact:** Ryan Roberts.

## Abstract

Osteosarcoma is an aggressive and deadly bone tumor, primarily afflicting children, adolescents, and young adults. Poor outcomes for osteosarcoma patients are intricately linked with the development of lung metastasis. While lung metastasis is responsible for nearly all deaths caused by osteosarcoma, identification of biologically defined, metastasis-targeting therapies remains elusive because the underlying cellular and molecular mechanisms that govern metastatic colonization of circulating tumor cells to the lung remains poorly understood. While thousands of tumor cells are released into circulation each day, very few can colonize the lung. Herein, using a combination of a novel organotypic metastasis *in vitro* model, single-cell RNA sequencing, human xenograft, and murine immunocompetent osteosarcoma models, we find that metastasis is initiated by a subpopulation of hypo-proliferative cells with the unique capacity to sustain production of metastasis promoting cytokines such as IL6 and CXCL8 in response to lung-epithelial derived IL1α. Critically, genomic and pharmacologic disruption of IL1 signaling in osteosarcoma cells significantly reduces metastatic progression. Collectively, our study supports that tumor-stromal interactions are important for metastasis, and suggests that metastatic competency is driven, in part, by the tumor cell’s ability to respond to the metastatic niche. Our findings support that disruption of tumor-stromal signaling is a promising therapeutic approach to disrupt metastasis progression.

## INTRODUCTION

Osteosarcoma is an aggressive bone tumor that primarily afflicts children, adolescents, and young adults and remains as deadly today as it was in the 1980s when cytotoxic chemotherapy regimens were first introduced. Outcomes for patients with osteosarcoma hinge almost exclusively on if patients develop lung metastasis. The 5-year overall survival rate for localized osteosarcoma is 70-75% and plummets to <20% for those that present with or subsequently develop lung metastasis[1]. Despite numerous clinical trials involving intensification of chemotherapy and addition of immunomodulatory agents, lung metastasis persists as a significant clinical challenge[2–5]. Novel therapies that disrupt the metastatic cascade could save up to 80% of lives currently lost to osteosarcoma[6–8].

Lung metastasis is a highly regulated and stereotyped cascade of events that involve invasion into surrounding stroma, intravasation into circulation, extravasation at distant sites, and successful colonization of the lung[9, 10]. Experimental and clinical evidence suggests that thousands of cells enter the blood stream daily[11]. Despite the deluge of tumor cells released into the blood stream, metastasis is an extremely inefficient process wherein <1% of tumor cells are capable of successfully colonizing distant organs[12, 13]. The discrepancy between circulating tumor cell density and development of metastasis underscores that potential metastatic sites are hostile environments and that tumor cells capable of colonizing and successfully surviving at metastatic sites are rare. Therefore, colonization is dependent on tumor education of the receiving microenvironment into a pro-tumoral niche[14]. How solid tumors such as osteosarcoma reprogram metastatic sites such as lung to promote metastasis remains poorly understood. Elucidating the mechanisms of osteosarcoma lung niche education will provide actionable vulnerabilities that prevent the emergence of metastasis.

Accumulating data in some cancers suggest that primary tumors are critical towards educating the secondary site into becoming more receptive to circulating tumor cells via formation of a pre-metastatic niche[15]. While there is significant data to support the importance of primary tumor conditioning of the pre-metastatic niche in some tumors through the release of cytokines, growth factors, or extracellular vesicles[16–18], it would be expected that patients with established primary tumors would most often present with clinically detectable metastatic disease. Indeed, tumors can efficiently re-recruit circulating cells through a process known as self-seeding[19]. In osteosarcoma, metastasis overwhelmingly occurs after the primary tumor has been resected[1]. The late emergence of metastasis after primary tumor resection suggests a complimentary hypothesis wherein subpopulations of circulating metastatically-competent cells can interact with and condition the lung into promoting metastatic growth.

Herein, we demonstrate that the subpopulation of cells which survive the metastatic bottleneck transform the metastatic lung niche into one that supports growth of recruited tumor cells. Mechanistically, osteosarcoma cells promote chemotaxis of other osteosarcoma cells through an IL6/CXCL8 based mechanism, two cytokines which are markedly upregulated by osteosarcoma-lung interactions and are critical for metastasis[20]. Utilizing single-cell RNA-sequencing (scRNA-seq) paired with ligand-receptor analysis, we uncover a novel mechanism whereby lung epithelial cells promote osteosarcoma IL6/CXCL8 via IL1. Epithelial induced IL6/CXCL8 production is limited to a small subpopulation of metastatic-competent cells that ‘anchor’ the lung metastatic niche and educate the niche towards a pro-metastasis state. From a translational perspective, we demonstrate that genomic and pharmacological disruption of IL1 signaling prevents osteosarcoma metastasis progression. Collectively, our study provides a therapeutically actionable mechanism of metastasis whereby specialized, metastatically competent tumor cells educate the metastatic niche through interactions with surrounding stromal cells.

## RESULTS

### Metastasis competent cells educate lung niche to promote metastatic colonization

Metastatic sites are inherently hostile towards disseminated tumor cells and therefore, must be transformed to accommodate metastatic colonization and growth. We hypothesized that osteosarcoma, which almost always targets the lung during metastasis, educates the lung through two possible mechanisms: 1) pre-metastatic niche conditioning by primary tumors or 2) disseminated cells with metastasis initiating capacity shape the metastatic soil into one that can support metastasis.

Utilizing a human xenograft experimental metastasis model (OS-17), we devised two experimental strategies to test how primary tumors influence lung metastasis (pre-metastatic niche conditioning). In the first experiment, which we called “Primary-No Primary,” we split mice into two groups, one in which osteosarcoma cells were orthotopically implanted into the tibia to form primary tibial tumors (Primary) and a second group which received a sham saline injection (No Primary). Once tumors in the Primary group reached a defined size of 1 cm^3^, both groups were intravenously injected with osteosarcoma cells and humanely euthanized 10 days after injection to quantify lung metastasis burden. We reasoned that if primary tumors conditioned the lung for metastatic colonization[21] and served as a source for circulating tumor cells, then the presence of a primary tumor would enhance metastatic seeding of circulating cells through both mechanisms. Surprisingly, we found that mice in the Primary group demonstrated significantly reduced metastatic burden compared to mice with no primary tumor (**Figure 1A**). We concluded that the presence of a primary tumor must influence circulating tumor cell dynamics by decreasing lung colonization.

**Figure 1.**
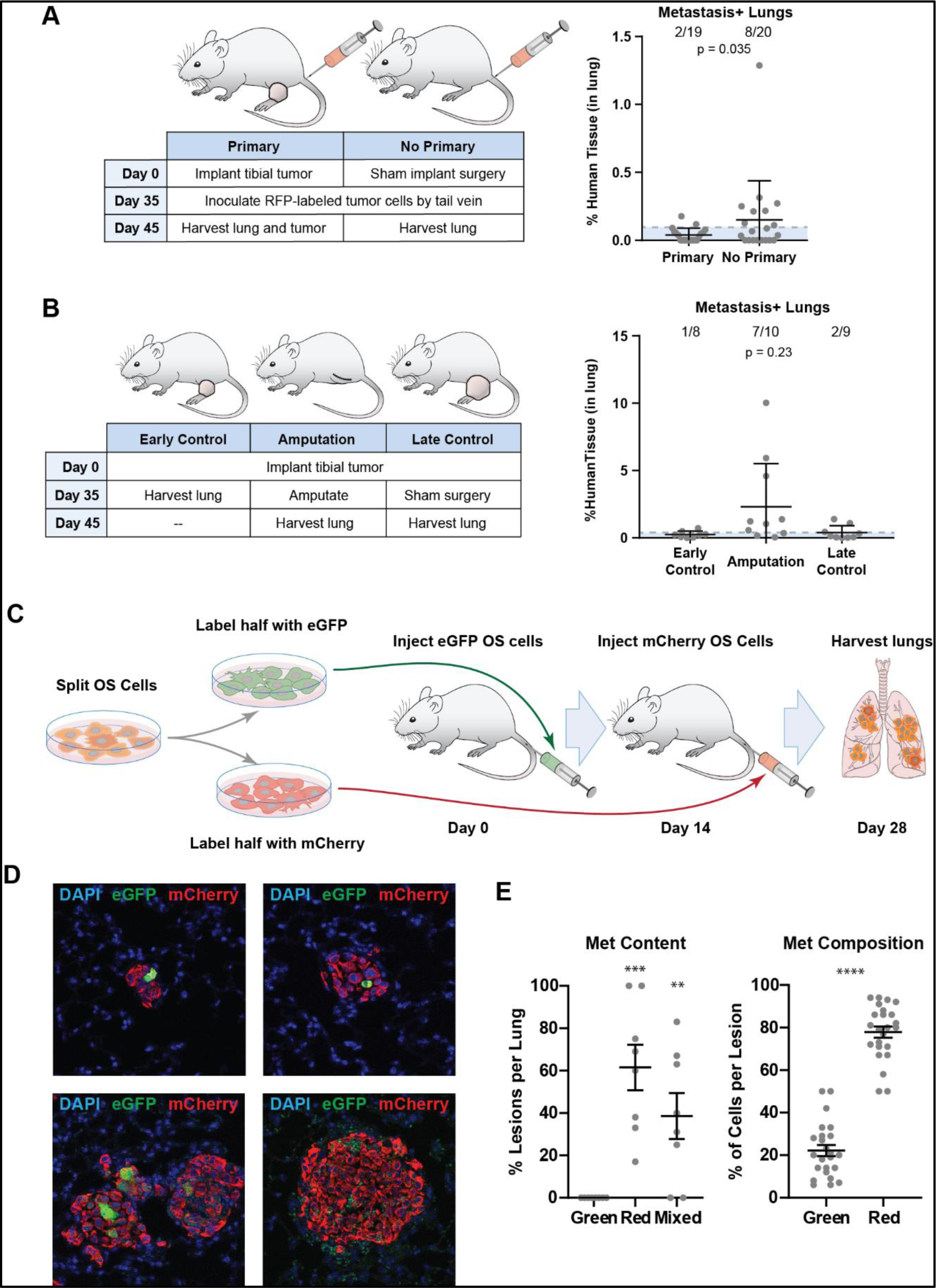
Metastasis competent cells educate lung niche to promote metastatic colonization. A) Schematic for ‘Primary-No Primary’ experiment with quantitation of metastasis burden by qRT-PCR. Fisher exact t-test, *p=0.035. B) Schematic for effect of surgical resection of primary tumor on lung metastasis with quantitation of metastasis burden by RT-PCR, Fisher exact t-test Amputation vs Early Control, p=0.23. C) Schematic of ‘Red versus Green’ experiment. D) Representative immunohistochemistry images of metastasis composition. E) Quantitation of metastasis content (all green versus all red versus mixed of both) and metastasis composition presented as percentage of each cell population in individual metastasis lesions. n=5 mice. ** p<0.01, *** p<0.001, **** p<0.0001 Welch’s t-test.

To complement the “Primary-No Primary” experiment, we next tested the effect of surgical resection of primary tumors on spontaneous lung metastasis. We randomized 30 mice into the following three cohorts (10 mice/cohort): The first cohort was euthanized at the time that the second group underwent amputation (35 days after tibial injection; Early Control), the second cohort had the affected leg amputated at day 35 post tumor implantation and were then euthanized on day 45 (Amputation). The last cohort was euthanized on day 45 post tibial injection, which represented the humane endpoint for mice bearing orthotopic tumors (Late Control). Lungs were harvested at the time of euthanasia and metastatic burden was quantified by qRT-PCR for human tissue. A combined 3 out of 20 mice in the Early and Late Control cohorts had detectable metastases, while 7 out of 10 mice undergoing amputation had detectable metastatic burden (**Figure 1B**). Thus, mice whose tumors were removed developed higher levels of tumor cell dissemination to the lung than mice that retained a rapidly growing primary tumor. These results hinted that the persistence of a primary tumor prevented either migration of tumor cells into the lung or growth of disseminated tumor cells rather than promoting lung metastasis.

We suspected that the mechanism explaining this observation might be a strong self-seeding effect, where disseminated tumor cells within the lung compete for circulating tumor cells with the large primary tumor[19, 22]. Therefore, we tested the alternative hypothesis that metastasis-competent cells shape the lung microenvironment towards one that supports and actively recruits circulating tumor cells. Lung metastasis is an inefficient process, even in experimental osteosarcoma metastasis models where tumor cells are injected directly into the bloodstream. In the OS-17 model, serial histological examination of lungs following intravenous tumor cell injection demonstrated that tumor burden drops dramatically from day 1 post-injection prior to nadiring at day 14 post-injection (**Supplementary Figure 1A-B**), consistent with a significant bottleneck following colonization and with previous data in other tumor models[12, 13]. We concluded that cells which persist after day 14 post injection represent metastasis-competent cells that ‘anchor’ the lung metastatic niche.

To test if these anchor cells create an environment conducive to recruiting/promoting tumor growth, we devised an experimental approach utilizing cells expressing green fluorescent protein (GFP) or mCherry (‘Red-green experiment’). This approach allowed us to distinguish anchor cells from cells recruited to established niche. First, mice were intravenously injected with OS-17-GFP cells. Fourteen days later, mice were injected with a second round of OS-17-mCherry cells. Mice were euthanized on day 28 after OS-17-GFP injection and processed for immunohistochemistry for GFP and mCherry respectively to quantify the percentage of metastases that were GFP, mCherry, or mixed (**Figure 1C-D**). Surprisingly, no identified lesion was composed of only GFP cells. mCherry+ cells colonized the regions surrounding GFP+ cells in significant numbers and constituted most of the cells in the mixed metastases (**Figure 1E**). These results suggested that anchor cells recruit circulating cells to established metastatic niches. This supported the conclusion that anchor cells educate the lung niche into a more favorable environment conducive to tumor growth. The ratio of mCherry-to-GFP cells also suggested to us that anchor cells may be hypo-proliferative given their relatively low density in mixed metastasis despite longer periods of potential growth, and that they could recruit other tumor cells with high proliferative capacity.

Collectively, these data argued that in osteosarcoma, the metastatic niche (at least in these models) is shaped more by the presence of metastasis-competent cells that “anchor” the lung niche than by the distant effects of primary tumors through a pre-metastatic niche conditioning mechanism. This experimental data fits with what is seen clinically where metastasis overwhelmingly occurs after primary tumor resection. We note that our data does not rule out a role for primary tumor-induced pre-metastatic niche conditioning in supporting metastasis, particularly in the colonization of anchor cells. We were intrigued by the idea that surviving disseminated tumor cells might influence circulating tumor cell dynamics. Our results suggested that osteosarcoma cells could promote chemotaxis of circulating tumor cells toward established niches—whether those be the primary tumor or a developing metastatic lesion. Therefore, we next sought to define the mechanisms by which anchor-type osteosarcoma cells promote the chemotaxis and recruitment of other osteosarcoma cells.

### Metastatic osteosarcoma cells promote osteosarcoma chemotaxis via IL6/CXCL8 mechanism

Our lab has previously demonstrated that inflammatory cytokines such as IL6 and CXCL8 play key roles in osteosarcoma metastasis[20, 23]. IL6 and CXCL8 are specifically over-expressed in metastasis relative to primary tumors and in cell lines with high-metastatic potential relative to cell lines with low-metastatic potential. Functional data supported that one role of these two pleiotropic cytokines was induction of cell migration[20]. Given our current data demonstrating tumor-induced recruitment *in vivo,* we hypothesized that the mechanism behind this recruitment might be IL6- and CXCL8-dependent chemotaxis.

We utilized reductionist *in vitro* migration assays to test this hypothesis. We first tested if cytokine-induced chemotaxis is correlated with metastatic potential. We constructed a panel of cell lines with high-metastatic potential (OS-17, 143B, MG63.3, LM7, K7M2 (murine)) and low metastatic potential (SaOS2, MG63, OS25) [24]. While all cell lines migrated to FBS regardless of metastatic potential (**Figure 2A and Supplementary Figure 2A**), only high-metastatic potential cell lines demonstrated marked cytokine-induced chemotaxis equal to or greater than FBS. Using the same cell line panel, we next tested if osteosarcoma cells induce chemotaxis of other osteosarcoma cells and if tumor-induced chemotaxis was IL6- and/or CXCL8-dependent (**Figure 2B and Supplementary Figure 2B**). Osteosarcoma-induced chemotaxis correlated with metastatic potential as evidenced by osteosarcoma conditioned media inducing cell migration that was equal to or greater than FBS alone in high-metastatic potential cell lines but not low-metastatic potential cells. In all cell lines except MG63.3, combined inhibition of IL6 (via GP130 inhibitor sc144) and CXCL8 (via CXCR1/2 inhibitor ladarixin) signaling resulted in a significant reduction in osteosarcoma-induced chemotaxis, largely reducing migration to at or below the unstimulated control. We concluded that osteosarcoma-induced chemotaxis is correlated with metastatic potential and requires IL6 and CXCL8. This data provided support for our working hypothesis that the observations made in **Figure 1** might be explained by tumor-tumor interactions that are mediated by IL6 and CXCL8 signaling.

**Figure 2.**
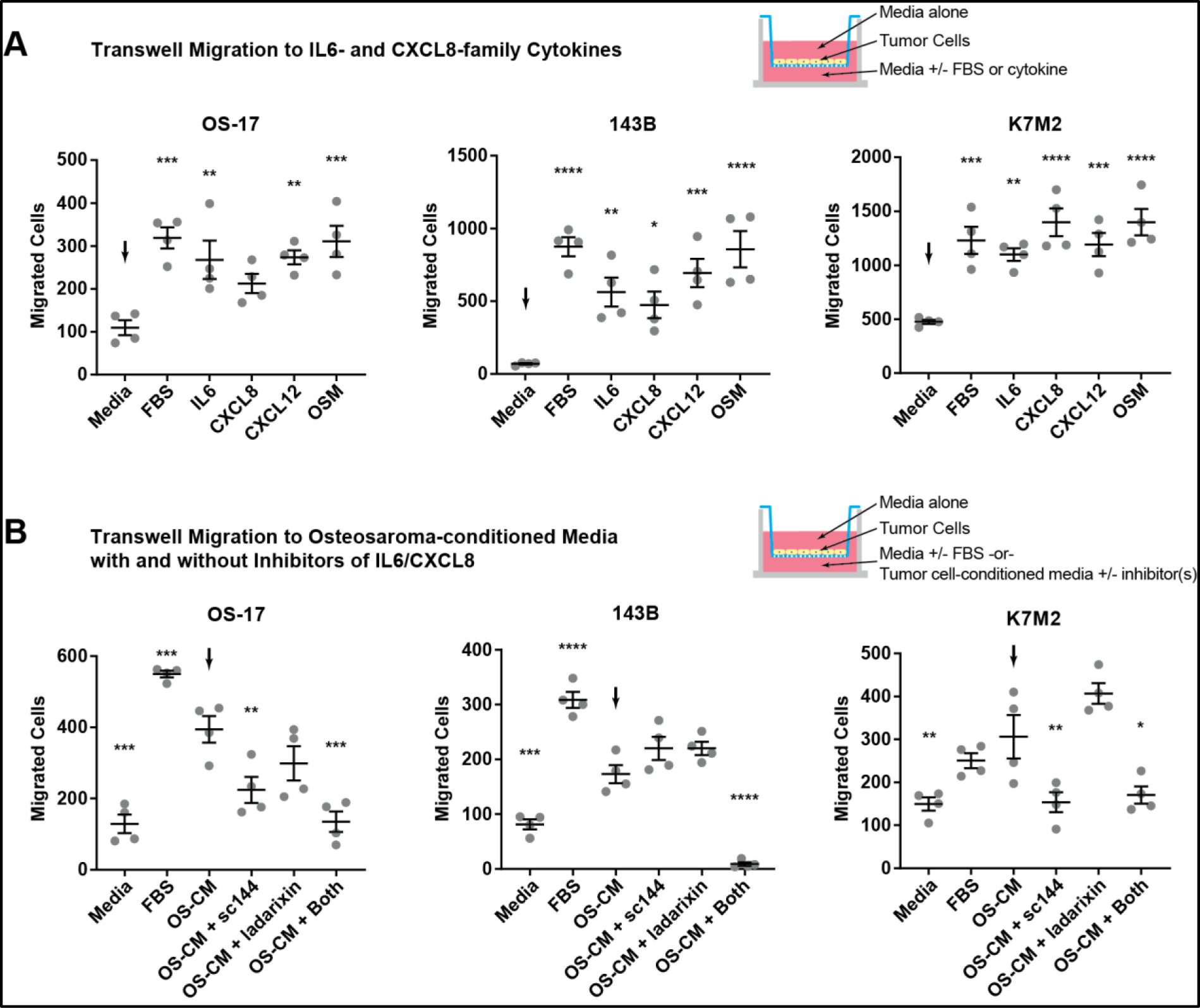
Metastatic osteosarcoma cells promote osteosarcoma chemotaxis via IL6/CXCL8 mechanism. A) Schematic for testing migration to IL6 and CXCL8 in transwell set-up and quantification of migrated cells relative to media control (down arrow). B) Schematic for testing role of IL6 and CXCL8 signaling in osteosarcoma-induced migration and quantification relative to vehicle treated osteosarcoma (OS) conditioned media (CM; arrow). **p=<0.01, *** p<0.001, ****p=<0.0001. with analysis of variance (ANOVA) followed by Tukey’s post hoc testing.

Indeed, we have previously demonstrated that IL6 and CXCL8 are markedly upregulated in metastatic tumors relative to their corresponding primary tumors and that metastatic osteosarcoma cells produce significantly more IL6 and CXCL8 in the presence of lung parenchymal cells [20]. If this were the mechanism active within our in vivo models, the production of IL6 and CXCL8 by metastasis-competent anchor cells within the early metastatic niche might be responsible for the accumulation of mCherry-labeled cells (the second injection) in the experiment shown in **Figure 1D**. However, the lung-derived signal initiating tumor cell cytokine secretion within the developing niche was not defined [20]. Therefore, we next sought to determine the mechanism by which tumor-lung interactions regulate IL6 and CXCL8 production by tumor cells.

### Epithelial derived IL1*α* drives tumor IL6/CXCL8 production

Osteosarcoma cells associate and make physical contact with lung epithelial cells upon lung colonization, which implicates them in directly influencing early osteosarcoma-lung niche dynamics, such as regulation of IL6/CXCL8[25]. Accordingly, we developed an *in vitro* co-culture system to allow for robust and unbiased evaluation of cell-cell communication networks between human lung epithelial cells and human osteosarcoma cells utilizing scRNA-seq. Interestingly, when OS-17 cells are cultured on top of a confluent layer of immortalized non-tumorigenic human bronchial epithelial cells (HBEC), they grow as spheroids that morphologically mimic early metastatic colonies *in vivo* (**Figure 3A**). HBEC monoculture, OS-17 monoculture, and HBEC:OS-17 co-cultures were dissociated and subjected scRNA-seq analysis. HBEC were identified by expression of cytokeratins (*KRT19*), while OS-17 cells were identified by expression of collagen (*COL1A1*, **Figure 3B**). Having separated tumor cells from lung epithelium and mono-from co-culture (**Figure 3C**), we next utilized NicheNet [26] to identify ligand:receptor (L:R) interactions that would be produced by HBEC (ligands), sensed by OS-17 cells (receptors), and drive the target gene changes observed in co-culture (including IL6 and CXCL8 production). *<u>Bona fide</u>* L:R interactions based on stringent selection criteria were identified (**Figure 3D**). IL1α and IL1β were notable for their strong regulatory potential of IL6 and CXCL8, which were among the top 50 most differentially expressed genes in co-culture compared to monoculture. (**Figure 3E-F**; arrows). We next sought to confirm these predicted ligand-receptor-target mechanisms that implicated epithelial IL1 in the regulation of osteosarcoma IL6 and CXCL8 production at the protein level.

**Figure 3.**
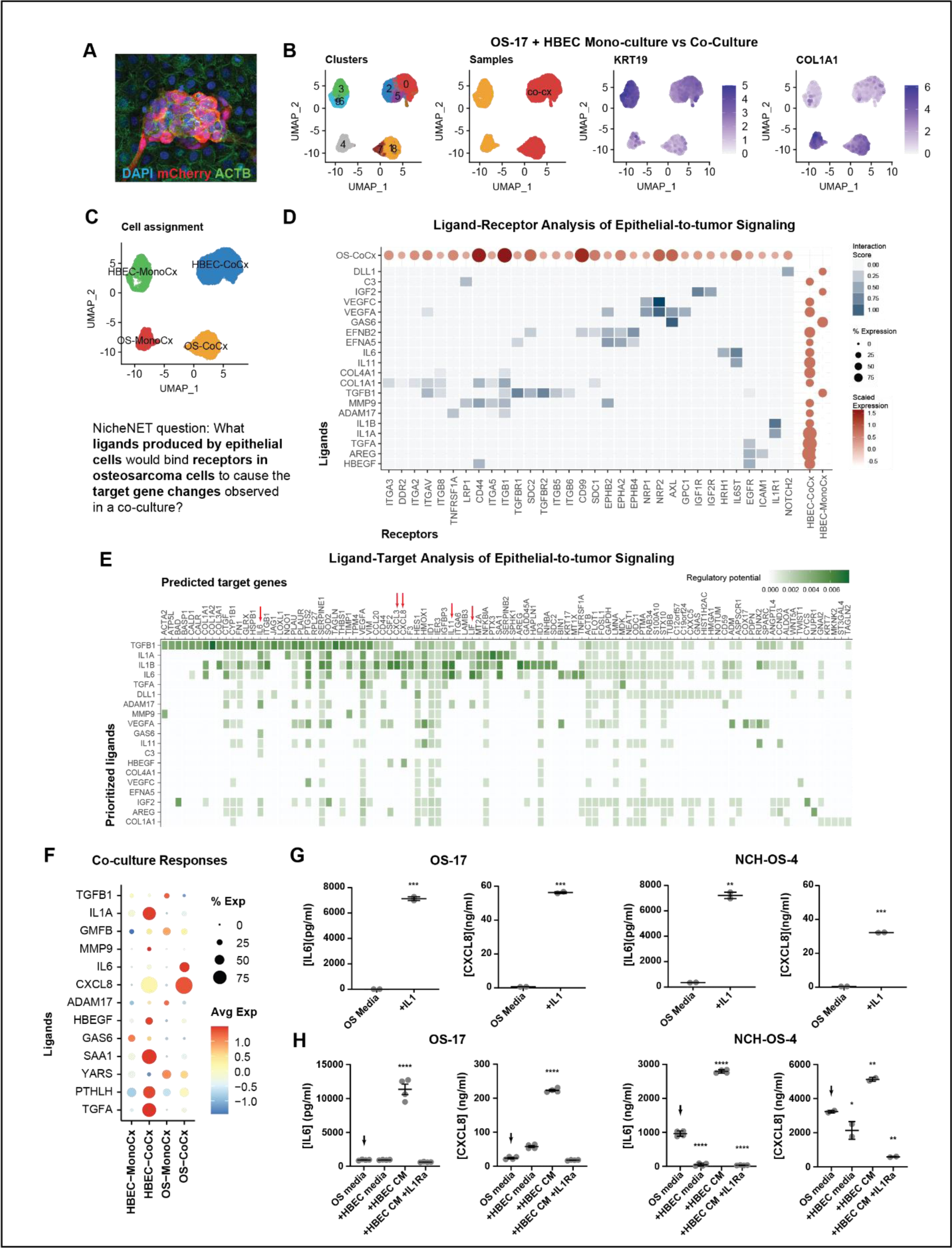
Epithelial derived IL1α drives tumor IL6/CXCL8 production. A) *In vitro* lung epithelial-osteosarcoma organotypic co-culture. Red marks mCherry expressing osteosarcoma cells, green=phalloidin (actin), and blue=DAPI. B-C) Single-cell RNA sequencing analysis of co-culture with Seurat based clustering on Uniform manifold projection (UMAP) with epithelial cells annotated by expression of *KRT19* and osteosarcoma cells annotated by expression of *COL1A1*. D) NicheNet heatmap and dotplot analysis demonstrating candidate epithelial-derived ligands (rows) osteosarcoma-derived receptors (columns). Strength of evidence for ligand-receptor interaction indicated by shade of blue in heatmap while the level of expression and percentage of cells expressing ligand or receptor are noted in dotplots. E) Heatmap demonstrating regulatory potential of epithelial-derived ligands for top 50 osteosarcoma differentially expressed genes (co-culture versus mono-culture). Red arrows note *IL6*, *CXCL3*, and *CXCL8*, which are strongly regulated *IL1* ligands. F) Dotplot of epithelial and epithelial (HBEC) ligands. Note IL6 upregulation in co-culture occurs in a minority of cells. G) IL1α significantly increases secretion of IL6 and CXCL8 in human osteosarcoma cells **p<0.01, ***p<0.001, ****p<0.0001 Welch’s t-test. H) Epithelial-induced (HBEC conditioned media (CM)) osteosarcoma IL6 and CXCL8 production requires IL1 signaling (IL1Ra= IL1 receptor antagonist anakinra). **p<0.01, ***p<0.001, ****p<0.0001. ANOVA followed by Tukey’s post hoc testing.

The IL-1 family of cytokines includes IL1α, IL1β, IL-18, and IL-33 and is widely implicated in inflammatory conditions and cancer[27]. ELISA demonstrated that both OS-17 and NCH-OS-4 (a metastatic, low passage patient-proximal cell line) cells stimulated with purified IL1 secreted significantly more IL6 and CXCL8 compared to unstimulated cells (**Figure 3G**). Having confirmed that both OS-17 and NCH-OS-4 cells were responsive to IL1 stimulation, we next confirmed that HBEC secreted the IL1 family of cytokines. IL1α but not IL1β was detected in HBEC supernatants under steady-state conditions (**Supplementary Figure 3A-B**). Interestingly, while co-culture itself did not elicit an increase in IL1α production, physical disruption of the epithelial cells drove a large increase in IL1α. This is interesting, given our recent report that lung colonization by osteosarcoma cells induces a pathological wound-healing program, which appears to be essential for lung colonization[24]. This may suggest that these two processes are, in fact, tightly linked. *In vivo,* IL1α demonstrated a membranous staining on peritumoral PDPN+ alveolar epithelial cells (AEC1). (**Supplementary Figure 3C**).

OS-17 cells and NCH-OS-4 cells were treated with conditioned media from HBEC (cHBEC) in the presence or absence of recombinant IL1 receptor antagonist (anakinra). In both cell lines, epithelial-induced IL6 and CXCL8 were significantly inhibited by anakinra (**Figure 3H**). Similar results were obtained when HBECs were replaced with small airway epithelial cell cultures (SAECs), which more precisely model the alveolar spaces of the lung (**Supplementary Figure 4**). Based on these data, we concluded that lung epithelial cells induce metastatic osteosarcoma cells to produce IL6 and CXCL8 in an IL1-dependent manner.

### Persistent IL6 and CXCL8 production is limited to a small subpopulation of cells

We have recently demonstrated that there is considerable intra-tumoral functional heterogeneity in osteosarcoma that is not attributable to changes at the genomic level [28, 29]. Indeed, the marked attrition of seemingly homogenous populations of highly metastatic cells noted upon lung colonization suggests significant functional heterogeneity regarding metastasis-generating capability. Based on the *in vivo* observations that IL6 and CXCL8 are required for metastasis, that only small subpopulations can survive the metastatic bottleneck and anchor the lung niche, and that only a fraction of the tumor cells appeared to produce IL6/CXCL8 in the co-culture experiments (**Figure 3F**) we hypothesized that IL6/CXCL8 expression is limited to a functionally defined subset of cells.

First, we queried our HBEC:OS-17 co-culture scRNA-seq dataset to explore heterogeneity with regard to IL6 and CXCL8 expression. Interestingly, after isolating OS-17 cells and performing a re-clustering analysis, we found that high expression of *IL6* and *CXCL8* was limited to a small cluster of cells, consistent with functional heterogeneity of IL6/CXCL8 expression (**Figure 4A**). We next stimulated OS-17 cells with IL1α for 72 hours (to mimic the time-course of the co-culture experiment) and assessed IL6 and CXCL8 heterogeneity at the protein level using immunofluorescence (**Figure 4B-C**). IL6 and CXCL8 expression was limited to approximately 20% of cells. We also assessed IL6 and CXCL8 protein expression in IL1α stimulated NCH-OS-4 and murine F420 osteosarcoma cells, which both demonstrated similar IL6 heterogeneity. The heterogeneous response to IL1α in two other cell lines ensured that our OS-17 observations were not cell line-dependent and likely represented a generalized response of metastatic osteosarcoma cells to IL1α. We next tested possible mechanisms that could explain the heterogeneity of IL1-induced IL6/CXCL8 response.

**Figure 4.**
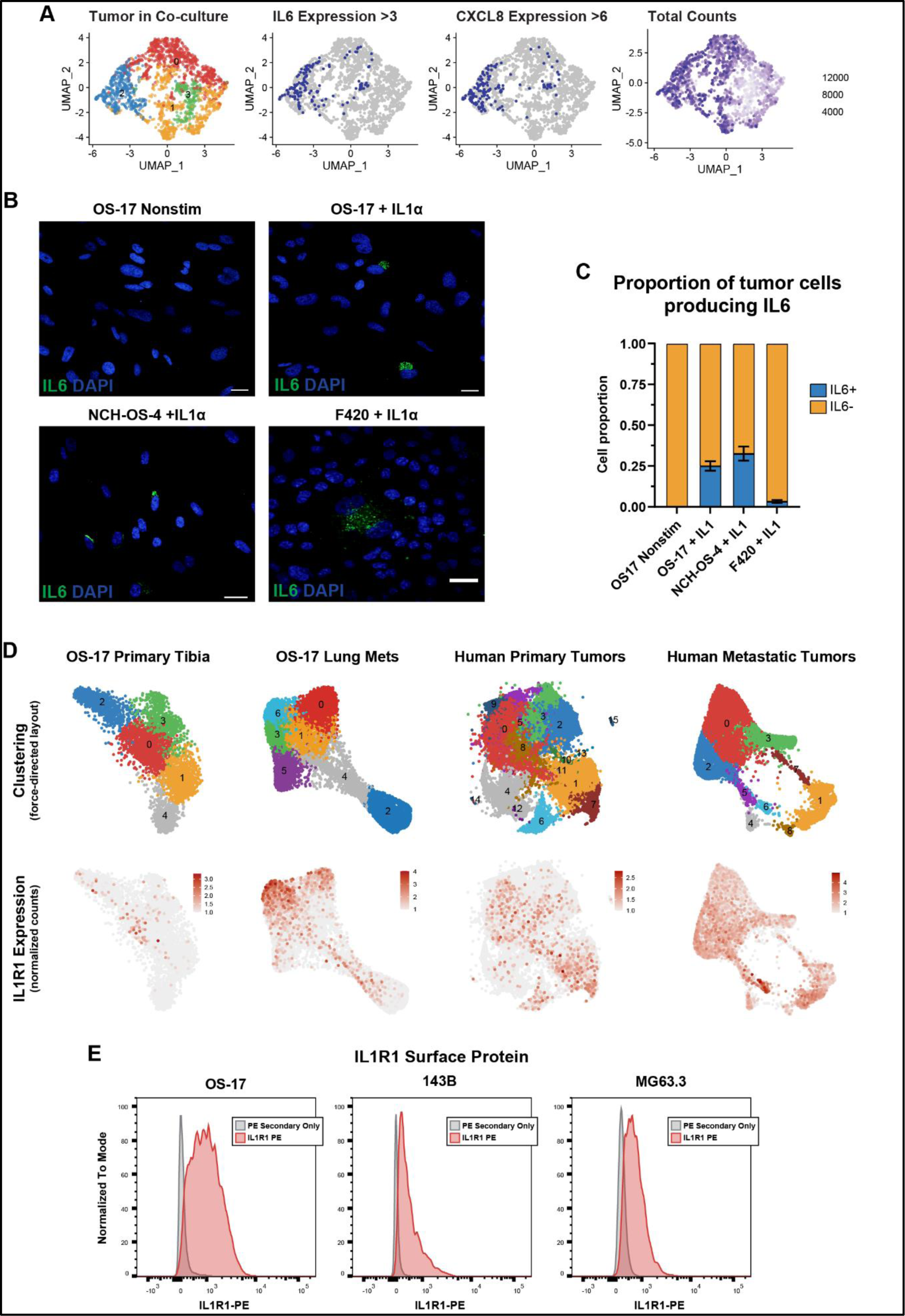
Persistent IL6 and CXCL8 production is limited to a small subpopulation of cells. A) UMAP of tumor cells subsetted from co-culture (Figure 3) demonstrating *IL6* and *CXCL8* expression is limited to a single cluster of cells. B-C) Immunofluorescence of osteosarcoma cells stimulated with IL1α (72 hours) and stained for IL6 (green) and DAPI (blue). Scale bar= 50µm. Percentage of IL6+ cells quantified in n=3 experiments, 100 cells/experiment. D) Single-cell RNA sequencing analysis of OS-17 tibial xenograft tumors, OS-17 experimental lung metastasis, human primary tumors, and human lung metastasis samples demonstrating heterogeneity of *IL1R1* expression. E) Flow cytometry analysis quantifying surface IL1R1 expression in human osteosarcoma cell lines.

Heterogenous expression of IL1R1 could explain cell-cell variation in IL6/CXCL8 output. To investigate this possibility, we isolated tumor cells from mice bearing OS-17 primary (orthotopic tibia) tumors or lung metastases and from a panel of human patient samples of both primary tumors and metastatic lesions, used those samples to generate 3’-capture single-cell RNA-seq libraries (10X Genomics), and evaluated expression of *IL1R1* across the tumor cell populations (**Figure 4D**). This analysis revealed marked heterogeneity of IL1R1 expression, with distinct subpopulations of cells showing significantly higher expression, mostly within the metastatic lesions. These findings were confirmed by flow cytometry, which suggested that the cell culture populations demonstrate a more homogenous IL1R1 expression by immunofluorescence at steady state (**Figure 4E**).

Furthermore, tumor cells demonstrated internalization of IL1R1 into punctate endocytic-like vesicles shortly after IL1α stimulation, consistent with active ligand-receptor binding (**Figure 5A**). Therefore, we concluded from these results that the heterogeneity of IL1R1 expression was not responsible for the heterogeneity of the cytokine-producing response *in vitro*. However, differences in IL1 signaling may be important for the emergence of anchor-type cells *in vivo*, especially within the metastatic niche.

**Figure 5.**
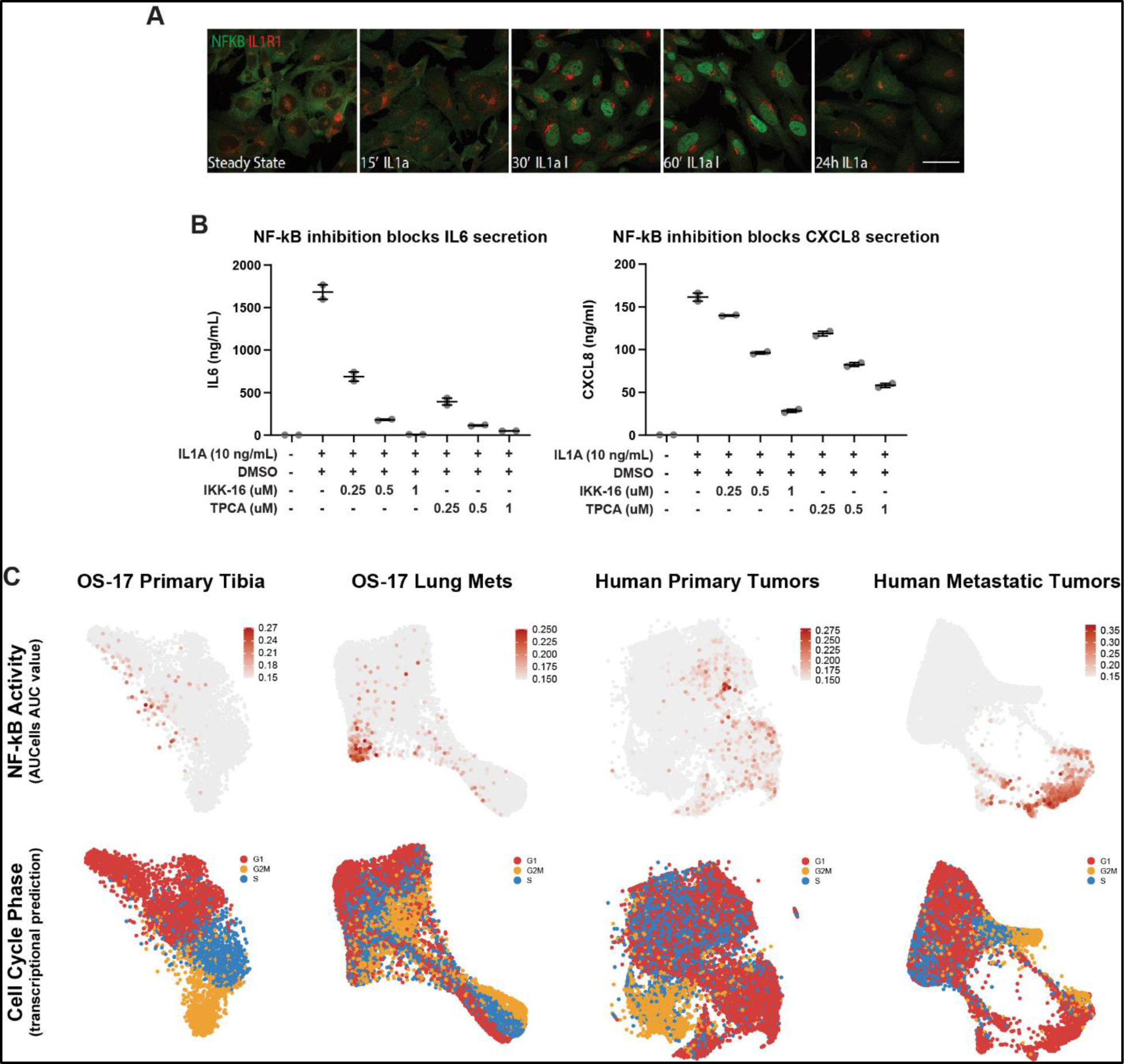
IL1α-induced IL6 and CXCL8 production requires NFkB signaling in a subpopulation of cells in G1 cell cycle phase. A) OS-17 cells were stimulation with IL1α, then fixed and stained for NFkB (green) or IL1R1 (red) at indicated timepoints. Note nuclear translocation of NFkB. Scale bar= 50µm. B) IKK inhibitors IKK-16 and TPCA-1 abrogate IL1α-induced IL6 and CXCL8 secretion as measured by ELISA. n=2 biological replicates done in technical triplicates. C) Single-cell RNA sequencing analysis of OS-17 tibial xenograft tumors, OS-17 experimental lung metastasis, human primary tumors, and human lung metastasis samples demonstrating restriction of NFkB activity (see methods) in small subpopulation of G1 cells.

Canonical IL1 signaling downstream of ligand:receptor binding leads to nuclear factor kappa B (NFκB) nuclear translocation[30]. Accordingly, we assessed if there was a heterogenous downstream signaling response to IL1α utilizing NFκB nuclear translocation as a marker. However, OS-17 cells generally demonstrated similar kinetics of NFκB translocation following IL1α. We next tested if IL1-induced IL6 and CXCL8 expression could occur through a NFκB-independent mechanism. To test the role of NFκB in IL1-induced IL6 and CXCL8 expression in OS-17 cells, cells were pre-treated with IKK-16 or TPCA, two distinct NFκB inhibitors, prior to IL1α stimulation, and IL6 CXCL8 secretion was then measured by ELISA (**Figure 5B**). In these experiments, IL1α-induced IL6 and CXCL8 secretion was effectively abrogated by NFκB inhibition. We therefore concluded that IL1-induced IL6 and CXCL8 requires NFκB pathway activation and that both heterogenous expression of IL1R1 and heterogenous activation of downstream signaling could play a role in the establishment of a hypersecretory subpopulation of tumor cells.

### Sustained IL6 production is maintained by a small subset of hypo-proliferative cells that anchor osteosarcoma cells to the lung niche

The nearly homogenous initial response to IL1α *in vitro* was intriguing in the setting of heterogenous IL6 and CXCL8 expression output late after stimulation. We performed an immunofluorescence time course for IL6 and CXCL8 expression to test if heterogeneity was defined by the persistence of IL6 and CXCL8 rather than initiation after IL1 expression stimulation. Strikingly, while nearly all cells demonstrate upregulation of IL6 and CXCL8 in the first 8-24 hours after stimulation, only 20% demonstrated persistence of IL6 and CXCL8 expression 72 hours post-stimulation (**Figure 6A**). Therefore, we concluded that sustained expression of IL6 and CXCL8 after IL1 is heterogeneous and confined to a small percentage of cells. We next sought to define further molecular characteristics of cells that persistently express IL6 and CXCL8 after IL1 stimulation.

**Figure 6.**
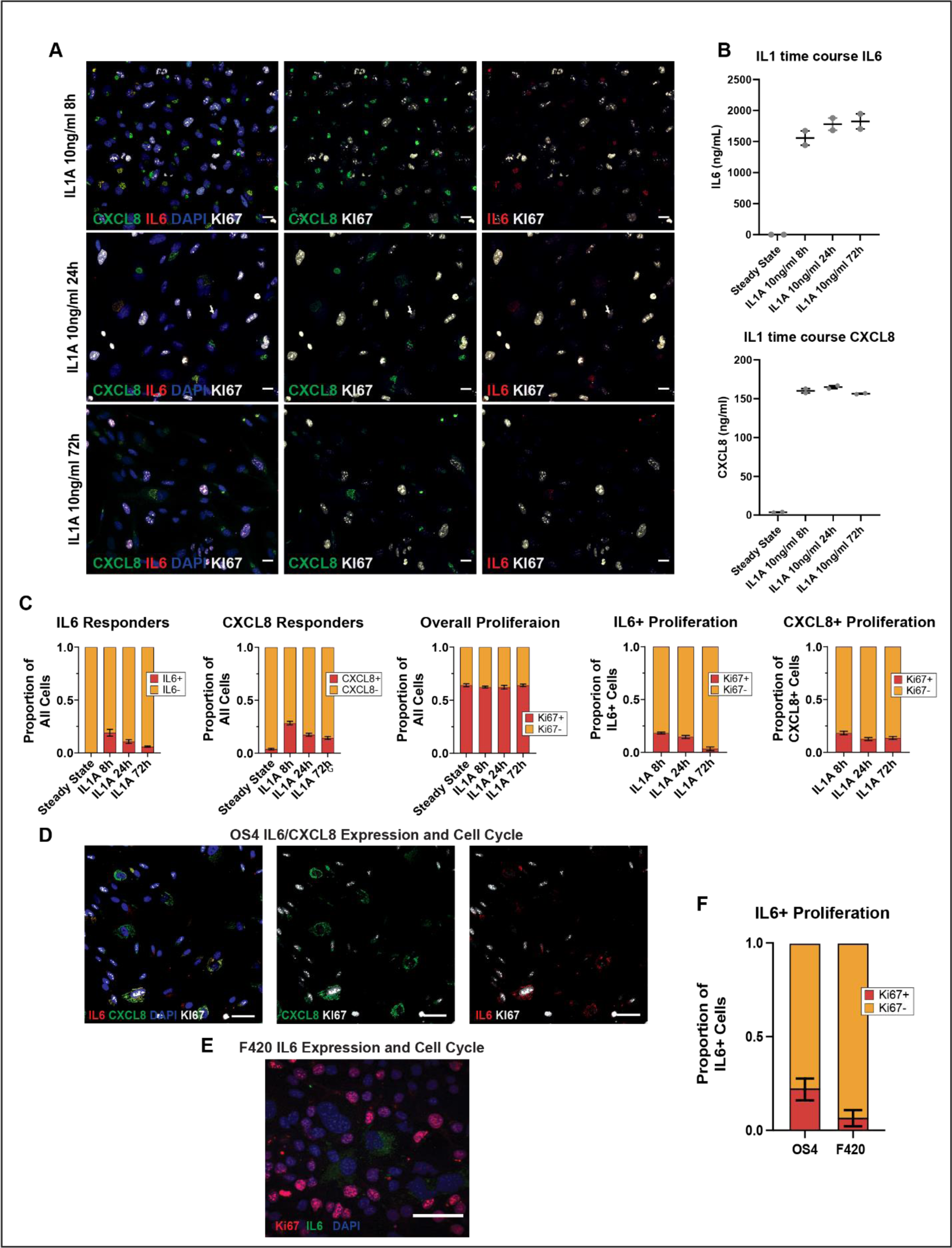
Sustained IL6 production is maintained by a small subset of hypo-proliferative cells that anchor osteosarcoma cells to the lung niche. A) Immunofluorescence expression of IL6(red) and CXCL8 (green) over time (8-72 hours) following stimulation with IL1α. White marks proliferating cells (Ki67). Blue=DAPI. B-C) ELISA and immunofluorescence quantitation demonstrating that IL6 and CXCL8 production remains constant over time while IL6 and CXCL8+ expression by immunofluorescence becomes progressively restricted to Ki67-cells. ELISA= 2 biological replicates done in technical triplicate. Percentage of IL6, CXCL8, and Ki67 quantified in n=3 experiments, 100 cells/experiment. D) IL6, CXCL8, and Ki67 expression in NCH OS-4 cells following stimulation with IL1α (72 hours). E) F420 murine cells stimulated with murine IL1α (72 hours) and stained with Ki67 (red), IL6 (green), and DAPI (blue). Scale bar= 50µm. F) quantitation of IL6 and Ki67 staining in NCH OS-4 and F420 cells.

Two hints as to the origins of this heterogeneity came from our previous experiments. First, the cells that survived the initial inoculation into mice (the GFP cells in the red-green experiment, **Figure 1D-E**) showed few signs of proliferation but significant evidence of tumor cell recruitment, which we subsequently showed was IL6/CXCL8-dependent (**Figure 2B**). Second, an overlay of cell cycle phasing onto the plots of NFκB pathway activation for a panel of *in vivo* osteosarcoma models and human patient samples (**Figure 5C**) suggests that the responding cell population remains largely within the G1 phase, especially in the metastatic lesions.

As cell cycle differences are a common source of heterogeneity within cell populations[31], we sought to validate whether cell cycle status changes in cytokine-producing cells. To confirm that persistent IL6-expressing cells represent a largely non-cycling population, we utilized multiparameter immunofluorescence to evaluate IL6, p21, and Ki67 expression simultaneously. OS-17 cells were stimulated with IL1α for 72 hours and processed for immunofluorescence to observe IL6, p21, and Ki67 (**Figure 6 A-C and Supplementary Figure 5A**). IL6-positive cells were significantly positive for the negative cell cycle regulator p21 and negative for the proliferation marker Ki67, suggestive of cells that were hypo-proliferative within the G1 phase. We confirmed that this was not limited to OS-17 cells as IL6 positivity was limited to Ki67-negative cells in our NCH-OS-4 cells and murine F420 osteosarcoma cells stimulated with human or murine IL1α, respectively (**Figure 6D-F**). We hypothesized that p21+/Ki67-cells may represent the IL6- and CXCL8-secreting anchor cells that survive the metastatic bottleneck *in vivo*.

The metastatic bottleneck represents a significant barrier for tumor cells to overcome to colonize distant organs, with few being able to withstand the selective pressure imposed during this process. We used immunohistochemistry to detect p21 and Ki67 in orthotopic OS-17 tibial tumors and during metastatic progression within the lung, reasoning that these markers would be enriched in early lesions (as the cells are able to respond to the lung microenvironment, i.e., produce IL6). p21 bright cells are rare in tibial tumors as well as end-stage metastatic lesions while making up the bulk of cells present two weeks after lung colonization (**Supplementary Figure 5B**), which corresponds to the period of the OS-17 bottleneck. As expected, p21 bright cells were negative for Ki67. Collectively, our *in vitro* and *in vivo* data supported a model that osteosarcoma lung colonization is mediated by a subpopulation of hypo-proliferative cells endowed with the capability of sustaining IL6 and CXCL8 production in response to lung niche signals (IL1α). To test the translational relevance of this model, we reasoned that inhibition of IL1 signaling would disrupt osteosarcoma lung metastasis.

### IL1 signaling is required for osteosarcoma metastasis progression

We first employed a genomic approach to assess the importance of IL1 signaling in osteosarcoma lung colonization. We utilized clustered regularly interspaced short palindromic repeats (CRISPR) technology to knockout (KO) IL1R1 in OS-17 cells. OS-17 cells were electroporated with a combination of plasmids, including cas9 and IL1R1-specific sgRNA, to disrupt IL1R1 expression. Successfully transfected cells depleted of IL1R1 were negatively selected utilizing fluorescence-activated cell sorting (FACS). Given our model, which supports that only a small subpopulation of cells is endowed with metastatic capability, we chose a negative selection FACS-based approach rather than isolating and expanding single-cell clones to select for IL1R1 KO cells to avoid clonal selection, which could confound the interpretation of results. Two independent IL1R1 KO cell lines did not produce IL6 or CXCL8 in response to cHBEC media relative to electroporation control cells, functionally validating IL1R1 disruption (**Figure 7A**). We next performed experimental metastasis assays with IL1R1 KO cells to assess the impact of IL1R1 KO on metastatic colonization *in vivo.* Experimental metastasis was induced via tail vein injection of OS-17 cells (OS-17-electroporation control, OS-17 IL1R1 KO CRISPR-B, or OS-17 IL1R1 KO CRISPR-C). Mice were monitored for signs of terminal disease, and once metastasis was confirmed in any mouse from any cohort, all mice were humanely euthanized and processed for histology to quantify metastatic burden. Three representative images from mice from each cell line are provided (**Figure 7B**). Mice injected with IL1R1 KO cells had significantly decreased metastatic lesions compared to mice injected with OS-17-electroporation controls (**Figure 7C**). We wondered if metastases that formed in IL1R1 KO mice represented subpopulations of OS-17 cells that were able to form metastases without functional IL1 signaling or if they represented small populations of OS-17 cells expressing IL1R1 that were not successfully selected against during our initial FACS procedure following CRISPR transduction. We performed IHC on electroporation control lungs and IL1R1 KO lungs to assess for IL1R1 expression and, interestingly, found that cells within the IL1R1 KO metastases expressed similar levels of IL1R1 found in control metastases (**Supplementary Figure 6E**). While this does represent a limitation to our study, as noted above, we reasoned that negatively selecting for IL1R1 expression by FACS would be less confounding compared to introducing possible clonal effects of growing out single cells. Therefore, our experiment inadvertently also showed a strong selection for the small populations of cells that escaped CRISPR-mediated IL1R1 KO, which also strongly supported a critical role for IL1 signaling in osteosarcoma metastasis.

**Figure 7.**
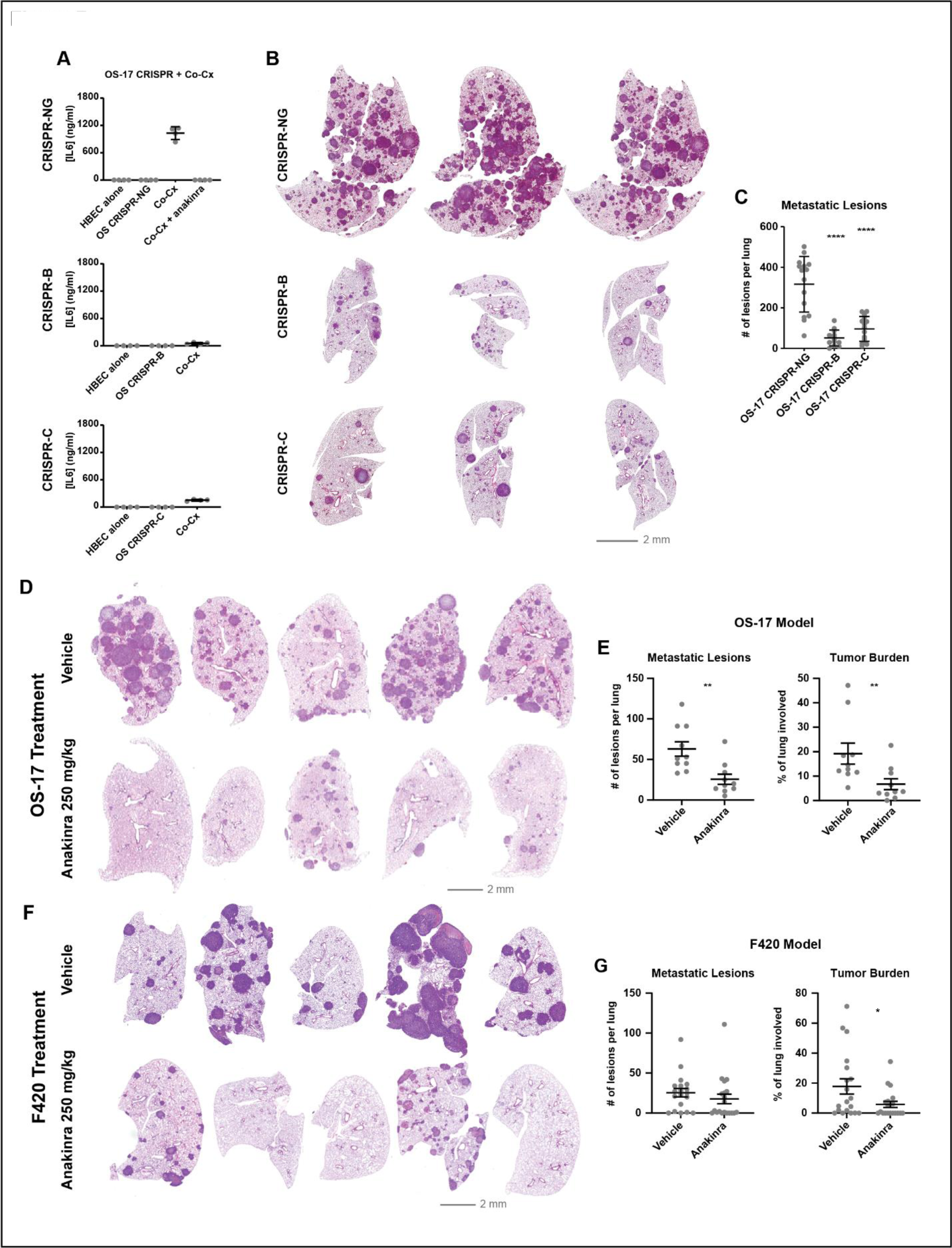
IL1 signaling is required for osteosarcoma metastasis progression. A) Functional validation of IL1R1 CRISPR knock-out by loss of epithelial-induced IL6 secretion measured by ELISA. n=4 biological replicates done in triplicate. B) Representative H/E stained lungs from mice inoculated OS-17 electroporated control (NG) or OS-17 IL1R1 CRISPR knock-out cell lines B and C. C) Number of metastatic lesions/ slice quantified by blinded examiner. n=15 mice/condition. **** p<0.0001. Welch’s t-test. D-E) Representative images and quantification of number of metastatic lesions and percent metastasis burden (relative to whole lung area) in mice inoculated via tail vein with OS-17 cells then treated with vehicle (PBS) or anakinra 1 day following tumor injection. **p<0.01. Welch’s t-test. F-G) Representative images and quantification of number of metastatic lesions and percent metastasis burden (relative to whole lung area) in mice inoculated via tail vein with F420 cells then treated with vehicle (PBS) or anakinra 1 day following tumor injection. *p<0.05. Welch’s t-test. Scale bar= 2mm.

To determine whether this pathway might represent a targetable vulnerability for developing metastatic lesions, we next utilized a pharmacological approach to disrupt IL1 signaling. Anakinra is a synthetic form of the IL1RA protein—a natural antagonist of IL1 signaling—with solid safety data and longstanding clinical experience, including in children and teenagers[32] with anakinra to strengthen the translational relevance of targeting IL1 in preventing osteosarcoma metastasis. Our initial attempts utilized dosing strategies based on published studies that have utilized anakinra in mouse inflammatory models[33] were not successful at preventing metastasis (25 mg/kg BID, **Supplementary Figure 6A-C**), nor at inhibiting IL1 signaling in the lung, as determined by pharmacodynamic analysis of phospho-IRAK4, a downstream signaling marker of IL1R1 activation[27] (**Supplementary Figure 6D**). Those pharmacodynamic studies did, however, show that higher doses of anakinra could sustain inhibition of the pathway in mice, so we proceeded to treat mice with high-dose anakinra (250mg/kg BID). Two days after tail vein injection, mice began treatment with subcutaneously injected anakinra or saline control twice per day until any mouse in either cohort showed clinical signs consistent with our pre-defined humane endpoints for lung metastasis. Upon confirming metastasis in one mouse, all mice were humanely euthanized, and lungs were harvested for histological analysis. High-dose anakinra significantly decreased the number of lesions as well as the overall metastasis burden within the lung in a dose-dependent fashion (**Figure 7D-E**). To increase rigor and assess the efficacy of anakinra in an immunocompetent model, we repeated the anakinra study using the F420 experimental metastasis model. Similar to OS-17, high-dose anakinra significantly disrupted F420 lung metastasis (**Figure 7F-G**). Collectively, we concluded that IL1 inhibition is a promising therapeutic strategy for preventing osteosarcoma metastasis progression.

## DISCUSSION

The development of lung metastasis, especially after primary tumor resection and chemotherapy, continues to lead to poor outcomes for young people diagnosed with osteosarcoma[1]. Clinical data demonstrating that metastasis overwhelmingly occurs after primary tumor resection has remained a fascinating yet perplexing phenomenon. In this study, we sought to devise experimental models that replicate this clinical phenomenon, which would allow us to elucidate factors that prevent the emergence of metastasis.

Utilizing a novel combinatorial approach of experimental *in vivo* platforms (Primary-No Primary, Early amputation, Red-green), we provide evidence to support that primary tumors influence circulating tumor cell fate and impair lung colonization, which agrees with recent findings from other groups[34, 35]. We significantly add to this growing body of literature, finding that in the presence of micro-metastatic disease, circulating tumor cells overwhelmingly favored recruitment to the established micro-metastatic lesions over initiating new ones. We concluded from this experiment that micro-metastatic lesions influence circulating tumor cell fate, supporting a model wherein cells which survive the stresses of the metastatic bottleneck initially ‘anchor’ in the lung and promote metastasis through at least two non-mutually exclusive mechanisms: directly recruiting circulating tumor cells through chemotaxis and indirectly by conditioning the hostile lung niche into one that promotes metastatic growth.

A logical progression from our initial data was to elucidate what distinguishes anchor cells that overcome metastatic bottleneck from the majority of osteosarcoma cells and how interactions with the surrounding niche influence anchor cell function. Notably, our study demonstrates that intra-tumoral heterogeneity and metastasis-initiating capacity are likely linked by the innate ability of metastasis-competent subpopulations to respond to the metastatic microenvironment. We found that anchor cells are defined by their ability to respond to lung epithelial derived IL1 and sustain production of two metastasis promoting cytokines IL6 and CXCL8. The exact functions of the pleotropic cytokines IL6 and CXCL8 in osteosarcoma metastasis remain enigmatic. However, our *in vivo* ‘Red-green’ experiment and *in vitro* functional migration data support that anchor cell IL6/CXCL8 promote the recruitment of tumor cells to the established metastatic niche. Our recent study elucidated that osteosarcoma metastasis progression is linked to fibrotic reprogramming of the lung niche[25]. As chronic inflammatory signaling is a well-known stimulus for fibrosis[36], epithelial-induced anchor cell derived IL6/CXCL8 may be critical for initiation of the metastasis associated fibrosis. This requires further study.

Because of the association of chronic inflammation with tumorigenesis, IL1 directed therapies are under intense clinical investigation in a variety of tumor types[37]. The interest in anti-IL1 therapies increased following results from the CANTOS trial wherein treatment of patients with atherosclerotic disease with IL1β antibody canakinumab markedly decreased the subsequent development of lung cancer relative to placebo-treated controls[38]. While cancer prevention was not the primary endpoint of the CANTOS study, it did provide rationale for dedicated canakinumab anti-cancer trials. Unfortunately, monotherapy or combinations of canakinumab with other agents have failed to statistically influence outcomes[39, 40]. While this may suggest anti IL1 therapy is more effective at preventing cancer than treating established cancer, it is important to note that canakinumab is directed at IL-1β. As our study demonstrated, IL1 signaling also occurs through IL1α, which is insensitive to canakinumab. Therefore, future studies with anakinra, a receptor antagonist, are warranted and are ongoing.

There are important limitations to consider with our study that will guide future areas of exploration. While we identified anchor cells as a small subpopulation of cells capable of colonizing and sustaining niche-induced inflammatory signaling required for metastasis progression, the underlying molecular mechanisms that regulate anchor cell identity remain incomplete. Our data suggests that these populations are hypo-proliferative, however, it is unclear whether they represent a terminally arrested population of cells or can re-enter cell cycle. From a translational perspective, our study supports the anti-metastatic effect of IL1 pathway inhibition but does not address when IL1 inhibition would be of most clinical benefit. Because IL1-driven inflammatory signaling initiates metastasis progression, and metastasis most often arises after primary tumor resection, the inclusion of anakinra in the adjuvant setting is reasonable but requires further study in orthotopic models combined with standard-of-care chemotherapy. Anakinra is FDA-approved and routinely used in children with inflammatory disorders[32]. The use of anakinra in the adjuvant setting is attractive given current agents used, such as receptor tyrosine kinase inhibitors like cabozantinib, which cause wounding healing issues following primary tumor resection[41].

In conclusion, our study supports a new model for osteosarcoma metastasis wherein a subpopulation of specialized tumor cells colonizes the lung and demonstrates how seemingly inert structural cells, such as epithelial cells, are critical in metastasis progression[42, 43]. Metastasis progression is initiated through lung epithelial-derived IL1 inducing osteosarcoma production of inflammatory cytokines such as IL6 and CXCL8. Lung metastasis progression is prevented by pharmacological inhibition of IL1 signaling with anakinra, suggesting that anakinra should be explored clinically.

## MATERIALS AND METHODS

### Cell culture

All cell lines utilized herein were routinely authenticated with STR and mycoplasma testing through Laboratory Corporation of America. Cells utilized for experiments were within 20 passage doublings after reviving from liquid nitrogen storage. OS-17 cells, a human osteosarcoma cell line, were derived from an early passage of the OS-17 patient-derived xenograft, grown from a primary femur biopsy at St. Jude’s Children’s Research Hospital in Memphis and was a gift from Dr. Peter Houghton. OS-25 cells were a gift from Øystein Fodstad at the Radium Hospital in Oslo. NCH-OS-4 cells, a human osteosarcoma cell line, were derived from an early passage of the NCH-OS-4 patient-derived xenograft, grown from a lung metastasis biopsy at Nationwide Children’s Hospital. SaOS2 cells were obtained from the American Type Culture Collection (ATCC, HTB85, RRID: CVCL_0548). LM7 cells were a gift from Dr. Eugenie Kleinerman from MD Anderson. All were maintained in RPMI (Corning, 10-040-CV) with 10% fetal bovine serum (FBS) (Atlanta Biologicals, S11150H). 143B, K7M2 and MG-63 cells were obtained from the American Type Culture Collection (ATCC, CRL8303, CRL2836 and CRL1427, RRID: CVCL_2270, CVCL_V455 & CVCL_0426). MG63.3 eGFP cells were a gift from Dr. Rosalind Kaplan at NCI. F420 cell were a gift from Dr. Jason Yustein at Baylor College of Medicine[44]. All were grown in DMEM (Corning, 10-013-CV) supplemented with 10% FBS. OS-17 cells were transduced using lentiviral particles designed to express green and red fluorescent protein (GFP, RFP/mCherry) and maintained in the same RPMI with 10% FBS as the parental OS-17 cells. HBEC3-KT cells were obtained from American Type Culture Collection (ATCC, CRL-4051) and maintained in Airway Epithelial Cell Basal Medium (ATCC, PCS-300-030) with the Bronchial Epithelial Cell Growth Kit (ATCC, PCS-300-040). HBEC:OS-17 organotypic co-culture were established as follows: HBEC were grown to confluence on 0.4µm inserts (Corning, 353090) with HBEC media in upper and lower chambers. OS-17 cells (50,000), were plated into top chamber in DMEM/F12 (Corning, 10-090-CV) + 1% FBS with HBEC media in lower chamber. Cultures were maintained until 72 hours prior to processing for single-cell RNA-sequencing (below).

### In vitro assays

For ELISA, OS cells were seeded into 24-well plates and incubated for 72 hours in basal medium alone or basal media supplemented with 10 ng/ml of either recombinant IL1α or IL1β (Biolegend, 570004 and 579404). Following incubation, cell-free supernatants were collected and analyzed by ELISA for changes in concentration of IL6 and CXCL8 using DuoSet ELISA Development Kits (R&D Systems, DY206, DY208). For imaging analysis, OS cells were seeded into 6 well plates that contained coverslips and incubated for 72 hours in basal medium alone, or basal media supplemented with 10 ng/ml of either recombinant IL1α or IL1β (Biolegend, 570004 and 579404) then fixed and processed as below.

#### Anakinra treatment

OS cells were seeded into 24 well plates and incubated for 24 hours in basal medium alone, or basal media supplemented with 100 ng anakinra per well. After 24 hours the cell media was aspirated and replaced with either basal medium alone, basal media + HBEC3-KT conditioned media (1:1) or basal media + HBEC3-KT conditioned media (1:1) plus 100 ng anakinra (supplied by Swedish Orphan Biovitrum AB, purchased from the inpatient pharmacy at Nationwide Children’s Hospital, Columbus, Ohio. NDC: 66658-234-01). After 72 hours cell-free supernatants were collected and analyzed by ELISA for changes in concentration of IL6 and CXCL8.

#### NFkB inhibition

Blockade of NFkB pathway was achieved by pre-treating cells with IKK inhibitors TPCA-1 (Selleckchem, S2824) or IKK-16 (Selleckchem, S2882) at indicated concentrations for 1 hour prior to IL1α stimulation. Cell free supernatants were collected at indicated timepoints and IL6 and CXCL8 concentrations determined as noted above.

### Production of lentiviral transduced cells

mCherry labelled OS-17 cells were synthesized using pLV-mCherry (Addgene, 36084) as described previously[45] with polyethylenimine (PEI) (Alfa Aesar, 43896) as the transfection reagent. OS-17 cells were transduced with in-house prepared mCherry virus or premade EF1A-GFP, CMV-RFP or CMV-Firefly luciferase lentiviruses (Cellomics, PLV-10018-50, PLV-10035-50 and PLV-10064-50) in the presence of Polybrene (8 μg/ml) (MilliporeSigma, TR1003G).

### Electroporation optimization of OS-17 cells

OS-17 cells were provided to the CRISPR/Gene editing core at the Research Institute at Nationwide Children’s Hospital (Columbus, OH). Cells were detached using TrypLE^TM^ (Thermo Scientific, 12605010) for ten minutes, then washed once with PBS. Once in suspension, 0.25×10^6^ cells were electroporated using two different electroporation kits and 4D-NucleofectorTM device (Lonza) to deliver 1 µg of pDNA encoding GFP. Sixteen hours after electroporation, cell populations were checked for GFP expressing live cells using fluorescent microscopy. OS-17 cells electroporated with SE buffer (Lonza) and the CM-104 program had the highest percentage of GFP+ cells, therefore these were the conditions used for subsequent CRISPR experiments.

### Generation of IL1R1-KO in OS-17 cells

To choose the best gRNAs for targeting of IL1R1 (Gene ID: 3554, Ensembl: ENSG00000115594), highest on-target and off-target scores were predicted by an algorithm using Benchling (Benchling.com). CRISPR-B gRNA sequence = TTTGTGTTGATGAATCCTGG. CRISPR-C gRNA sequence=AAACTACCCGTTGCAGGAGA. Synthetic gRNA (Synthego), HiFiCas9/RNP complexes were suspended in SE electroporation buffer (Lonza) and electroporated using the 4D-Nucleofector system (Lonza) program CM-104 in the presence of electroporation enhancer (IDT). Wildtype OS-17 cells were also electroporated without Cas9/RNP as a negative control. Following electroporation, cells were kept in basal media (RPMI supplemented with 10% FBS). IL6 ELISA was used to functionally IL1R1 knockout.

### Migration Assay

Transwell migration assays were performed using 24 well plates and transwell inserts (Falcon, 353097). OS cells (1×10^4^) were plated into the inserts and cells were exposed to appropriate chemoattractant factors. After 24 hours, the inserts were removed, media was drained, and cells on the upper surface of the membranes were removed with a cotton swab. Each insert was stained using the Quik Dif staining kit (Siemens, B4132-1A) and allowed to dry per manufacturer guidelines. Images were obtained using an inverted microscope, and cell numbers were counted using Adobe Photoshop’s counting tool. For experiments involving conditioned media, OS cells were grown in their appropriate media for 72 hours; media was collected, spun down to remove cellular debris, and added to the bottom chamber. For experiments using serum as a chemoattractant, top chambers contained RPMI only, while bottom chambers contained 1% serum. For experiments that used cytokines as a chemoattractant, a concentration of 50 ng/ml of each was used (BioLegend, 570804, 574204, 581202, and 555202). In experiments testing the ability of small molecules to block serum-induced migration/invasion, 10 μM sc144 (a small molecule inhibitor of IL6ST, the signaling receptor for IL6) (Sigma, SML0763) and/or 100 nM DF2156A (ladarixin, a small molecule inhibitor of CXCR1 and 2, the receptors for CXCL8, kindly provided by Dompé Farmaceutici, L’Aquila, Italy) was added to the media.

### Human IL1R1 Flow cytometry

OS cell lines from tissue culture were evaluated for IL1R1 expression using flow cytometry (BD LSRFortessa, BD Biosciences). Cells were grown under their standard cell culture conditions stated above and harvested at 1×10^6^ cells/100 µL in PBS for IL1R1-specific staining. Cells were incubated with blocking IgG at 1 µg/10^6^ cells (Normal Goat IgG, sc-2028, Santa Cruz Biotechnology, Inc.) for 15 minutes at room temperature. After blocking incubation, excess blocking IgG was not removed, and cells were stained for IL1R1 (IL1R1 PE-conjugated, R&D Systems, FAB269P) or control antibody (Goat IgG Control PE-conjugated, R&D Systems, IC108P) at 10 µL/10^6^ cells for 30 minutes at room temperature in the dark. Next, stained cells were washed 3 times with MACS Buffer (1:20 in PBS, Miltenyi Biotec, 130-091-376) to remove any remaining unbound antibody. Cells were centrifuged at 500 x g for 5 min after each wash for a total of three washes with MACS buffer. Finally, stained OS cells were resuspended in 400 µL of MACS buffer for flow cytometric analysis.

### qPCR evaluation of human metastatic tissue in mouse lungs

qPCR assays were developed to determine the quantity of human tissue compared to mouse tissue in human osteosarcoma metastasis-bearing mouse lungs. Primers targeting each gene were designed using NCBI Primer-BLAST[46] to meet the following criteria: amplicon smaller than 125 bp, pair must be separated by at least 1 intron (where possible), preference for 5′ mRNA targets, no nonsynonymous target recognition for potential amplicons smaller than 1,500 bp, and no potential off-species amplification. Primer pairs were validated using cDNA extracted from murine and human cell lines. Amplification was performed TaqMan™ Universal PCR Master Mix (Applied Biosystems) on an Applied Biosystems HT 7900 thermocycler. Accepted primer pairs demonstrated efficiency greater than 92% (as calculated using “window of linearity” methods [47]), single, narrow curves on melting curve analysis, and consistency across all 4 validation specimens. Sequences for each of the primer pairs are listed here: XenoH-F9-Probe 5’-/56-FAM/CAT CTT CCT /ZEN/CAA ATT TGG ATC TGG CTA TGT AAG TGG CTG G/3IABkFQ/ −3’, XenoH-F9-Forward 5’-TTG CCC TTC TGG AAC TGG AC −3’, XenoH-F9-Reverse 5’-GAA GAC ATG TGG CTC GGT CA −3’ and XenoM-F9-Probe 5’-/56-FAM/TTC CAC TGG /ZEN/TGG ATA GAG CCA CAT GCC TTA GGT /3IABkFQ/-3’, XenoM-F9-Forward 5’-TCA GTG GCT GGG GAA AAG TC −3’, XenoM-F9-Reverse 5’-GGGTCC CCC ACT ATC TCC TT −3’. cDNAs obtained from murine lungs from our primary vs. no primary and the amputation study (see below) were then analyzed by qPCR using each primer set separately, run in triplicate. Dissociation curves were run for every assay well. Less than 5% of wells were censored from further analysis due to signs of nonspecific amplification in the dissociation curves. Cycle threshold (Ct) values were determined visually for each primer set using Applied Biosystems SDS software. Relative quantification was determined and then normalized to expression within the primary tumor for each primer set.

### *In vivo* osteosarcoma studies

All animal experiments were carried out in accordance with the Institutional Animal Care and Use Committee (IACUC, protocols AR15-00022 and AR14-00045) of the Research Institute at Nationwide Children’s Hospital (Columbus, OH) guidelines. The approved protocols aimed to minimize the number of mice used and to minimize any pain or distress. All manipulations occurred under sterile conditions in a laminar flow hood. Staff performed all surgical procedures under general anesthesia and aseptic conditions. Mice used were 6-8 weeks-old CB17SC scid-/- (RRID: IMSR_TAC:cb17sc) female mice for the OS-17 models, C57Bl/6 (RRID: MGI:2159769) for the F420 model. Mice were maintained under barrier conditions.

#### ‘Primary-No Primary’

A murine model of circulating tumor cells was developed and used to evaluate the influence of the presence or absence of a primary appendicular OS on the seeding of the circulating tumor cells. Under general anesthesia and surgical asepsis, a per-cutaneous intra-tibial injection was performed in two groups of mice. Group 1 received an injection of 5×10^5^ GFP-labeled OS-17 cells through the tibial plateau. Group 2 received a sham injection of RPMI, without tumor cells. In group 1, tumors were measured once weekly, and mice were weighed twice weekly and examined for signs of illness or swelling around the surgical site. Timepoint 1 was determined to be when tumors grew to 1 cm^3^. As mice from group 2 did not have a tumor growing, they were paired with mice from group 1. At this time, mice from both groups received a tail vein injection of 1×10^6^ RFP-labelled OS-17 cells. Ten days after tail vein injection was defined as the end point. At endpoint, each mouse was euthanized and primary tumor (if present), blood, bone marrow and lungs were collected. The percentage of human tissue (or OS cells) in the lungs was quantified using RT-PCR as noted above.

#### Effect of primary tumor resection on metastasis

An orthotopic murine model of appendicular OS was developed to investigate the influence of the surgical removal of the primary tumor on the development of metastasis. Under general anesthesia and surgical asepsis, a per-cutaneous intra-tibial injection was performed with 5×10^5^ GFP-labelled OS-17 cells. Tumors were measured once weekly, and mice were weighed twice weekly and examined for signs of illness or swelling around the surgical site. Timepoint 1 was determined to be when tumors grew to 1 cm^3^. Timepoint 2 (endpoint) was 10 days after time point 1. Once mice had reached criteria for time point 1, they were randomly assigned to the 3 study groups. The control group (Early Control) was euthanized at timepoint 1 (tumor volume 1 cm^3^), and organs were harvested. The treatment group (Amputation) underwent amputation of the tumor-bearing limb by coxo-femoral disarticulation at time point 1 and was euthanized at time point 2. The no treatment group (Late Control) did not undergo amputation and were euthanized at timepoint 2. Blood, bone marrow and lungs were collected at endpoint. The percentage of human tissue (or OS cells) in the lungs was quantified using RT-PCR (as noted above).

#### ‘Red-green experiment’

An experimental murine model of osteosarcoma metastasis was developed to distinguish between recruited circulating tumor cells to an established niche from metastasis-initiating ‘anchor’ cells. At day 0, mice were tail vein inoculated with 1×10^6 GFP-labelled OS-17 cells. Fourteen days later, mice were tail vein inoculated with 1×10^6 mCherry-labelled OS-17 cells. After 28 days from the initial GFP-labeled OS-17 inoculation, mice were euthanized, and their lungs were collected for immunohistochemical examination of lungs for GFP/mCherry composition of metastases as noted below.

#### Experimental metastasis progression

Mice were tail vein inoculated with 1×10^6 OS-17-luciferase tagged cells. At various time points (day 0, 1, 7, and 14), luciferase-based imaging was performed to non-invasively assess the presence of lung metastases. At each time point, two mice were euthanized, and lungs were collected and evaluated for metastatic burden.

#### Effect of IL1R1 knockout on metastasis

Mice were tail vein inoculated with 1×10^6^ OS-17 IL1R1 knockout cells (CRISPR-B or CRISPR-C) or OS-17 electroporated control cells (CRISPR-NG). Mice were continuously monitored for signs of metastatic endpoint, and once a mouse from any group was confirmed to bear metastatic disease upon necropsy, all mice were humanely euthanized. At the conclusion of the study, lungs were collected and evaluated for metastatic burden via histological examination.

#### In vivo anakinra treatment

Mice were inoculated with OS cells (1×10^6^ for OS-17 and 5×10^5^ for F420) via tail vein injection. Mice were then divided into two treatment groups. The first treatment group received a dose 250 mg/kg Anakinra via subcutaneous injection, twice daily. The second treatment group received sham injections (PBS) for the duration of the study. Mice were observed as above until any mouse from that group reached endpoint, at which time all mice from that group were euthanized. Endpoint criteria was defined as a Body Condition Score (BCS) greater than 3, or loss of greater than 10% body mass in a single mouse. At the conclusion of the study mice were euthanized and lungs were collected and evaluated for metastatic burden. Anakinra (Lot. 33719-1A) was supplied by Swedish Orphan Biovitrum AB, purchased from the inpatient pharmacy at Nationwide Children’s Hospital, Columbus, Ohio. NDC: 66658-234-01.

### Tissue/cell dissociation, single cell RNA library preparation, sequencing, and analysis

Lungs harvested from mice were processed using the human tumor dissociation kit (Miltenyi Biotec, 130-095-929) with a GentleMacs Octo Dissociator with Heaters (Miltenyi Biotec, 130-096-427). HBEC mono-cultures, OS-17 monocultures, or HBEC:OS-17 co-cultures (72 hours) were gently trypsinized and dissociated into single cell suspensions. Following dissociation into single cell suspension, single-cell 3’ RNA-Seq libraries were prepared using Chromium Single Cell 3′RNA-sequencing system (10X Genomics) with the Reagent Kit v3.1 (10X Genomics, PN-1000121) according to the manufacturer’s instructions. cDNA library quality and quantity were determined using an Agilent 2100 bioanalyzer using the High Sensitivity DNA Reagents kit (Agilent, 5607-4626) and then sequenced on an Illumina NovaSeq 6000. Raw data and initial processing was performed using cellranger version 7 (10X Genomics) with reads mapped to hg19 version of the human genome.

Raw data and initial processing was performed using Cell Ranger version 7.2 (10X Genomics) with reads mapped to the mm10 version of the mouse genome (mouse tumor samples), the hg38 version of the human genome (human patient samples), or a mixed genome containing both (xenograft samples). Mapped and aligned cell by count matrices were loaded into Seurat toolkit version V5[48] for further analysis. Cells with <200 detected genes or mitochondrial DNA >20% were removed. RNA counts were normalized with regularized negative binomial regression via SCTransform. After dimensionality reduction with principal component analysis and UMAP embedding, unsupervised clustering was achieved using “FindNeighbors” and “FindClusters” functions in Seurat with dimensions 1:30 and resolution of 0.3 clustering of all cells and 0.6 for subclustering. Automated cell type annotation was performed using SingleR[49] using the Human Primary Cell Atlas and Tabula Muris as annotation references. Differential gene expression between clusters was determined using “FindAllMarkers” function with parameters of only.pos=TRUE, min.pct=0.25, and logfc.threshold=2. Clusters with less than 50 cells or less than 20 differentially expressed genes were removed. Cell clusters were annotated based on canonical marker expression. Tumor cells were identified based on the expression of classical osteosarcoma genes (COL1A1, COL1A2, SATB2, RUNX2, FOSL1) and by their poor matching to the normal reference standards in the automated cell type annotation. Candidate ligand (epithelial)-receptor(osteosarcoma) interactions driving differential gene expression were identified using NicheNet.

### Gene set enrichment analysis and pathway graph-based projection

In this study, we considered genes to be differentially expressed if they demonstrated log2FC >2 or <-2 changes in gene expression and the adjusted p-value was <0.01. Differentially expressed gene tables were assessed with GSEA (ClusterProfiler[50]) against Hallmark and KEGG pathways (msigdbr[51]). Subset analysis of candidate pathways (such as NFkB signaling) was performed using AUCell to determine the area-under-the-curve for expression of genes within the designated pathway (in this case, the “TNFA signaling via NFkB” pathway from the Hallmark compilation (msigdbr). AUC scores were merged into the Seurat metadata for visualization.

### Histology

Sternotomy was performed and lungs were flushed via slow instillation of phosphate buffered saline (PBS) (Corning, 21-031-CV) through right ventricle until lungs were cleared of blood. Lungs were dissected from mouse *en bloc* along with trachea and heart. Lungs were slowly instilled with 10% neutral buffered formalin (NBF) (Sigma-Aldrich, HT5012-1CS) until fully expanded, then trachea was tied off with suture and placed in NBF for 24 hours at 4°C. After repeated washing in PBS, lung lobes were carefully dissected from each other and processed for paraffin embedding (FFPE) and immunohistochemistry (IHC). For FFPE, 4µm sections were deparaffinized with xylene and rehydrated through ethanol series. To quantify metastasis burden, FFPE sections were counterstained with hematoxylin and eosin via conventional methods. Heat-mediated antigen retrieval methods utilized citrate buffer (CB; pH 6.9) or Tris-EDTA buffer (TE; pH 9) with Tween 20 (Fisher Bioreagents, BP337-100) at 0.2% v/v. Antigen retrieval (AR) buffer used was antibody dependent (see below).

### Immunohistochemistry (IHC), Immunofluorescence (IF), and microscopy

Cells growing on #1.5 glass coverslips (Electron Microscopy Sciences, 72204-04) were fixed in NBF for 10 minutes at room temperature. Following three rinses in PBS, coverslip or histological sections were then incubated for at least 1 hour in blocking solution consisting of PBS, 0.2% triton100 (v/v) (Sigma, X-100-100ml), and 3% bovine serum albumin (w/v) (Sigma, A-7888) at room temperature. FFPE sections were rehydrated in PBS for 30 minutes then blocked as above. Primary antibodies used: rabbit anti-CXCL8 (Abcam, ab289967, IF), mouse anti-human vimentin (Abnova, SRL33, IF, AR=TE), rabbit anti-p21 (Cell Signaling Technology (CST), 2947, IF, AR=CB), rabbit anti-NFkB (CST, 8242, IF) hamster anti-Podoplanin (Developmental Studies Hybridoma Bank(DSHB), 8.1.1, AR=TE), rat anti-Ki67 (Invitrogen, 14-5698-82, IF, AR=CB), mouse anti-human IL 6 (R&D, MAB2061, IF), mouse anti-human IL1α membrane form 488 (R&D, FAB2001G, IF), goat anti-mouse IL1α (R&D, AF-400-NA, IF, AR=TE). Primary antibodies were diluted in blocking solution and sections were incubated in primary antibody solution overnight at 4°C. Following three rinses in PBS, sections were incubated with appropriate AlexaFluor labeled secondary antibodies (Invitrogen), DAPI (Invitrogen, D1306), and where indicated, Phalloidin AlexaFluor 647 (Invitrogen, A12379) or Wheat Germ Agglutinin (WGA) AlexaFluor 647 (Invitrogen, W11261) diluted in blocking solution for 1 hour at room temperature. Following PBS rinses, coverslips and tissue sections were mounted in Fluoromount G (Invitrogen, 00495802).

### Microscopy and image analysis

Fluorescent microscopy images were obtained on a LSM 800 confocal microscope using 20X air or 63X oil objectives. Tile-scanning was used for high-resolution imaging of large areas with images stitched together with 10% overlap using ZenBlue software. CZI files were then uploaded and processed using ImageJ. All manipulation of images (brightness/contrast and smoothing) were done uniformly to each image. The cell counter plugin was used to manually quantify cell populations by IL6, CXCL8, p21, or Ki67 positivity. Metastasis burden was quantified from H/E sections by blinded examiner using Aperio ImageScope Software (V12.4.3). Whole lung lobe images were obtained with Aperio FL ScanScope digital slide scanner with 20X objective.

### Statistical analysis

Statistical analysis was performed using Prism V9. Number of samples and mice are explicitly stated in text of results or figure legends with exact p-values. Data reported with error bars represent mean +/- standard error of the mean (SEM).

## Data sharing and availability

Code used to analyze single cell RNA-seq (scRNA-seq) data can be found on GitHub at github.com/kidcancerlab. All single-cell RNA data Will be available via GEO.

