## Supplemental Figure 1 for "Metastasis-initiating osteosarcoma subpopulations establish paracrine interactions with both lung and tumor cells to create a metastatic niche"

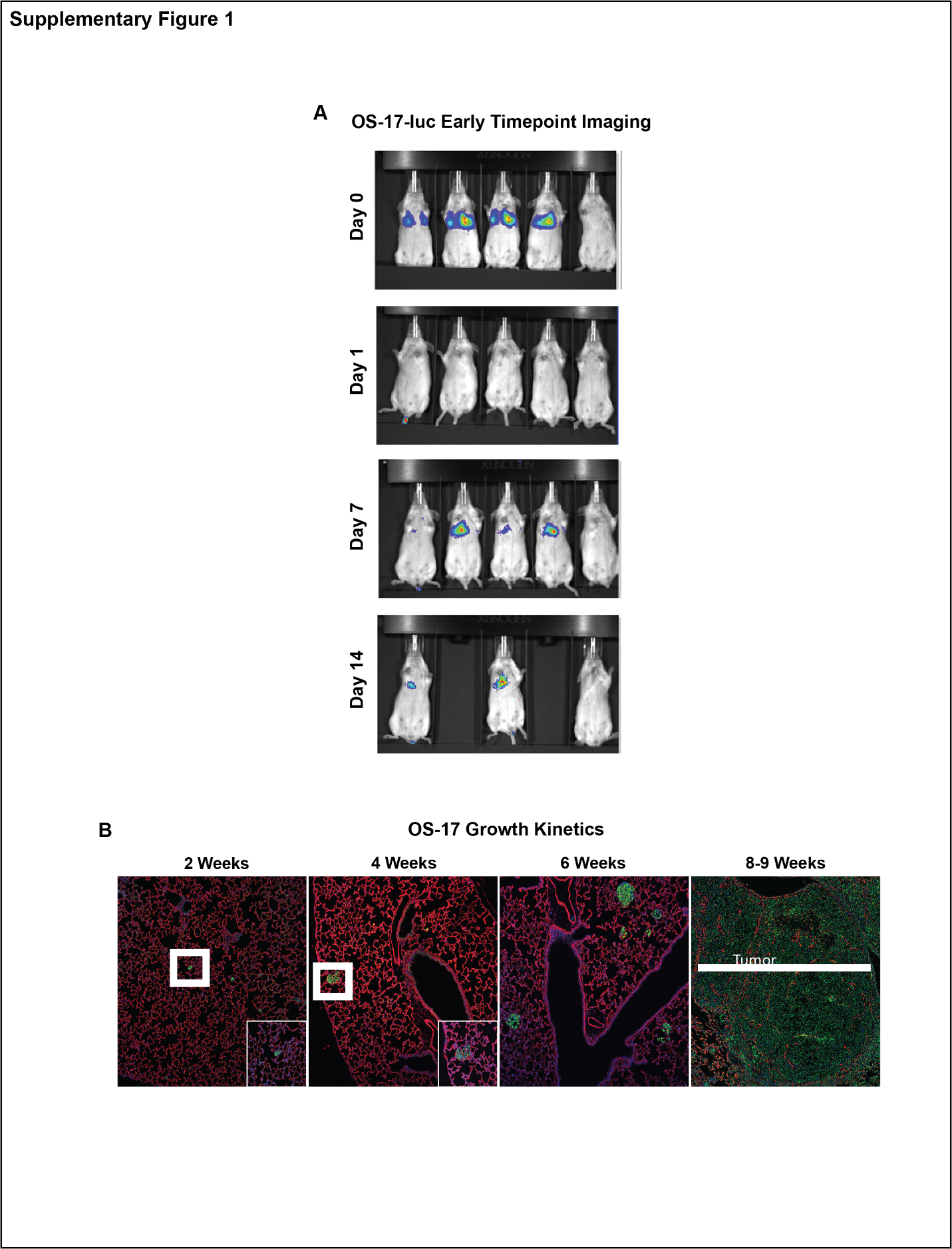

**Supplementary Figure 1.** A) Luciferase based monitoring of metastasis progression in OS-17 xenograft model. Note marked attrition of cells between day 0 and day 14. B) Immunohistochemistry of experimental metastasis progression in OS-17 model. White box denotes inset. Red= Wheat germ agglutinin. Green= human vimentin (marks tumor cells).
