## Supplemental Figure 2 for "Metastasis-initiating osteosarcoma subpopulations establish paracrine interactions with both lung and tumor cells to create a metastatic niche"

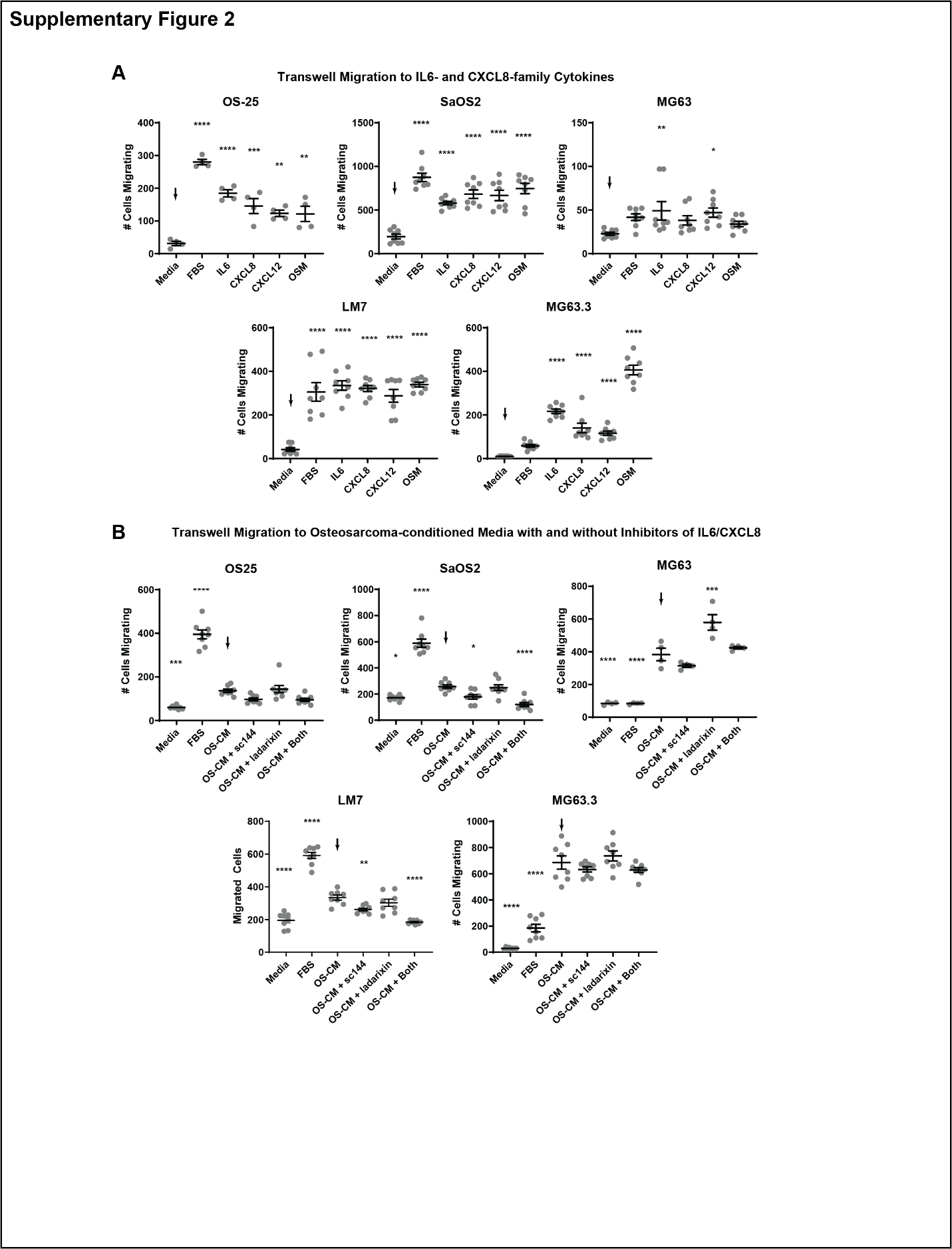

**Supplementary figure 2**. A-B) Extended panel of cell lines measuring migration in response to IL6 , CXCL8, and osteosarcoma conditioned media (OS-CM) in presence or absence of IL6 and/or CXCL8 inhibition. OS-25 SaOS2 and MG63 all have low metastatic potential *in vivo* while LM7 and MG63.3 are highly-metastatic derivatives of SaOS2 and MG63 respectively.. ****p<0.0001 ANOVA followed by Tukey’s post hoc testing.
