## Supplemental Figure 3 for "Metastasis-initiating osteosarcoma subpopulations establish paracrine interactions with both lung and tumor cells to create a metastatic niche"

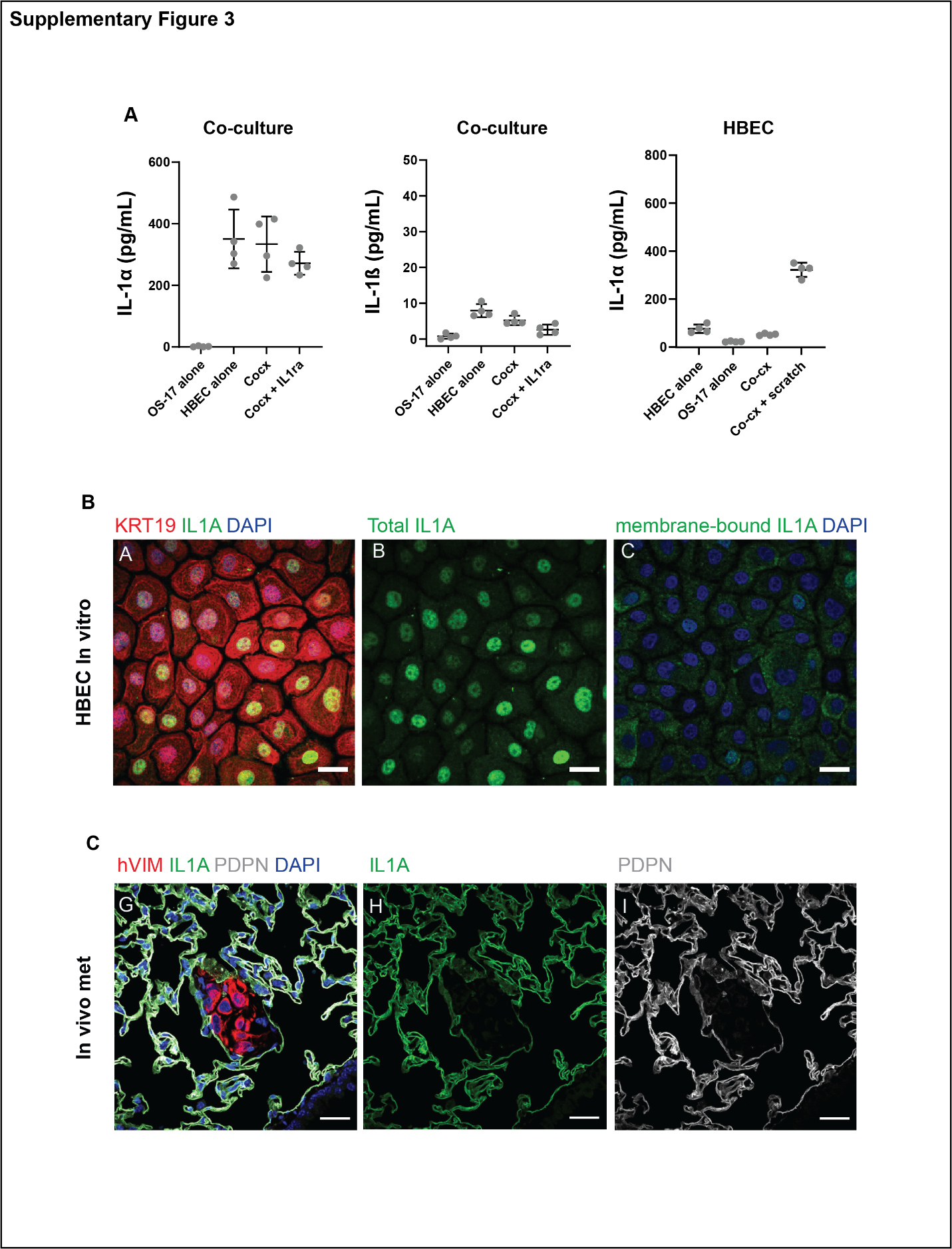

**Supplementary figure 3.** Epithelial cells express and secrete IL1α *in vitro* and *in vivo*. ELISA measuring IL1α and IL1β in osteosarcoma cells and epithelial cells or in co-culture in the presence or absence of anakinra. n=4 biological replicates done in technical triplicates. B) Immunofluorescence of total IL1α and membrane-bound IL1α in HBEC in culture. Red=KRT19, green=IL1α, blue=DAPI. C) IL1α (green) is expressed on the membrane of peri-metastatic (red) PDPN-expressing type 1 alveolar epithelial cells (white). Blue=DAPI. Scale bar= 50µm.
