## Supplemental Figure 4 for "Metastasis-initiating osteosarcoma subpopulations establish paracrine interactions with both lung and tumor cells to create a metastatic niche"

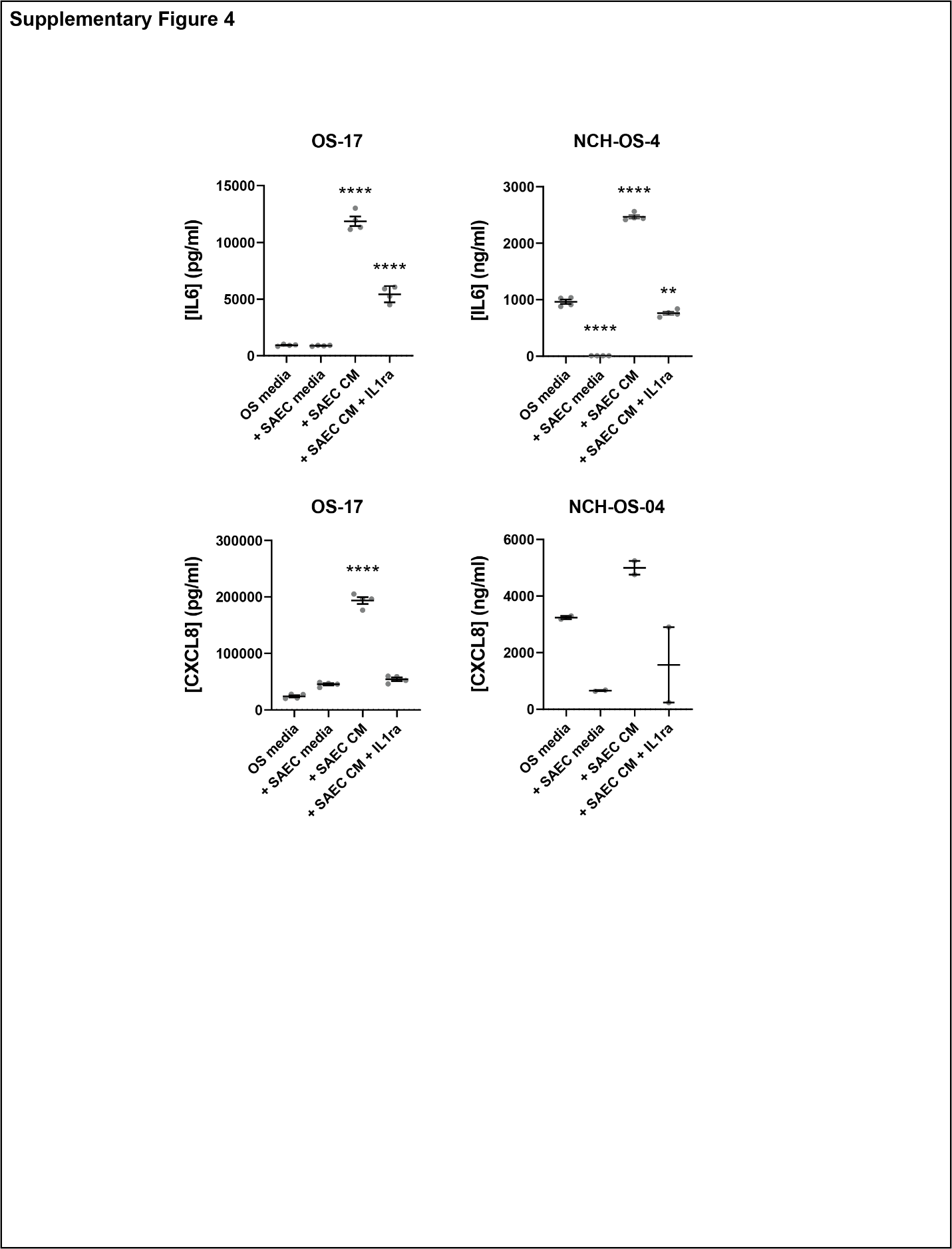

**Supplementary figure 4**. Small airway epithelial cells (SAEC) also stimulate IL6 and CXCL8 production in OS-17 and NCH OS-4 cells in an IL1-dependent manner as measured by ELISA. **p<0.01, ****p<0.0001, ANOVA followed by Tukey’s post hoc testing. n=4 biological replicates done in technical triplicates.
