## Supplemental Figure 5 for "Metastasis-initiating osteosarcoma subpopulations establish paracrine interactions with both lung and tumor cells to create a metastatic niche"

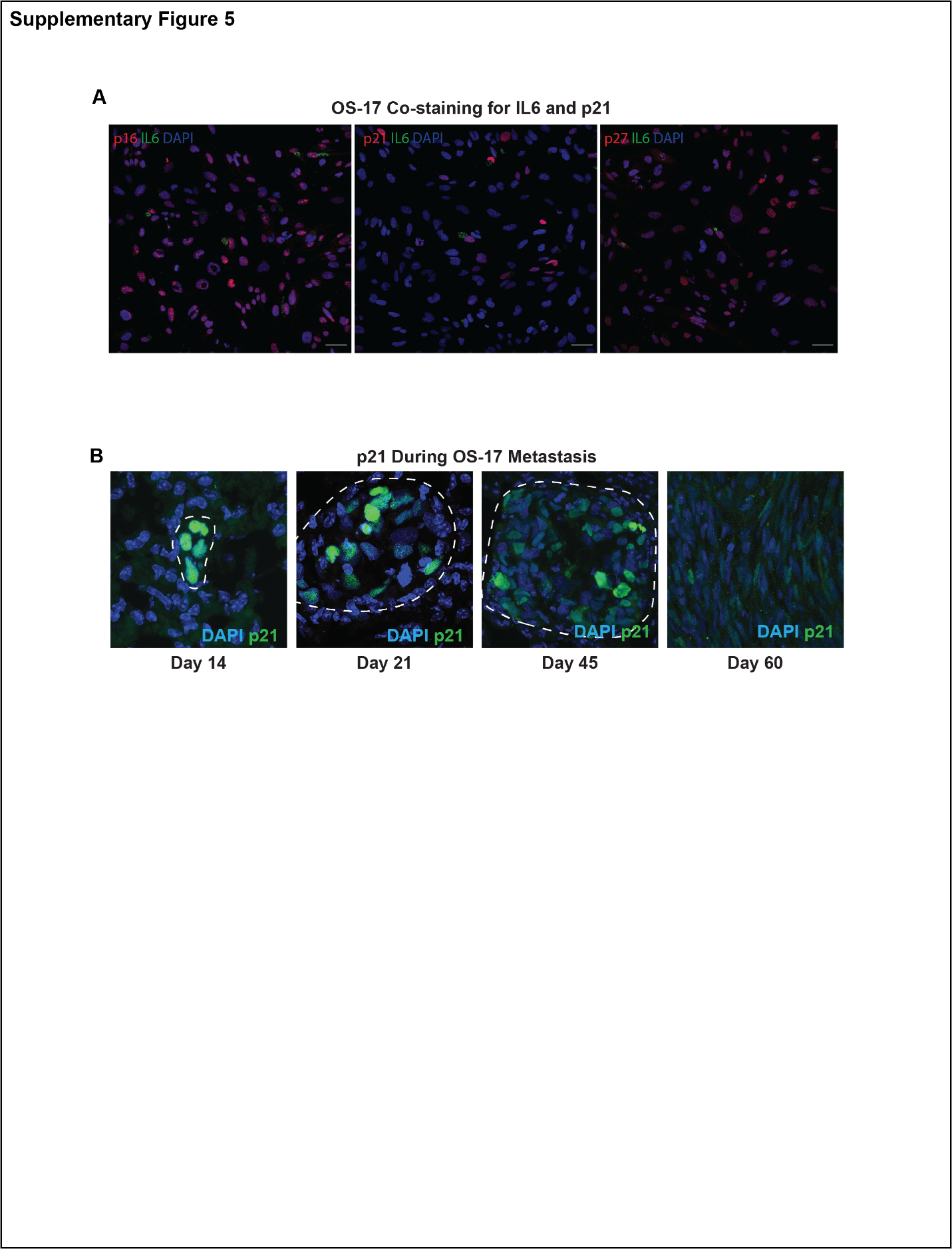

**Supplementary figure 5**. IL6 producing anchor cells expressed p21 *in vitro* and p21+ cells survive metastatic bottleneck *in vivo*. A) OS-17 cells were stimulated with IL1α for 72 hours then fixed and stained for negative cell cycle regulators p16, p21, and p27 (red) and IL6 (green). IL6+ cells are p21+. B) OS-17 cells were inoculated into lung and mice euthanized at indicated time points and lung processed for p21 immunohistochemistry. Early metastatic cells (14 days) are p21+ while established metastases (Day 60) have very few p21+ cells. Green=p21, Blue=DAPI. Scale bar= 50µm.
