## Supplemental Figure 6 for "Metastasis-initiating osteosarcoma subpopulations establish paracrine interactions with both lung and tumor cells to create a metastatic niche"

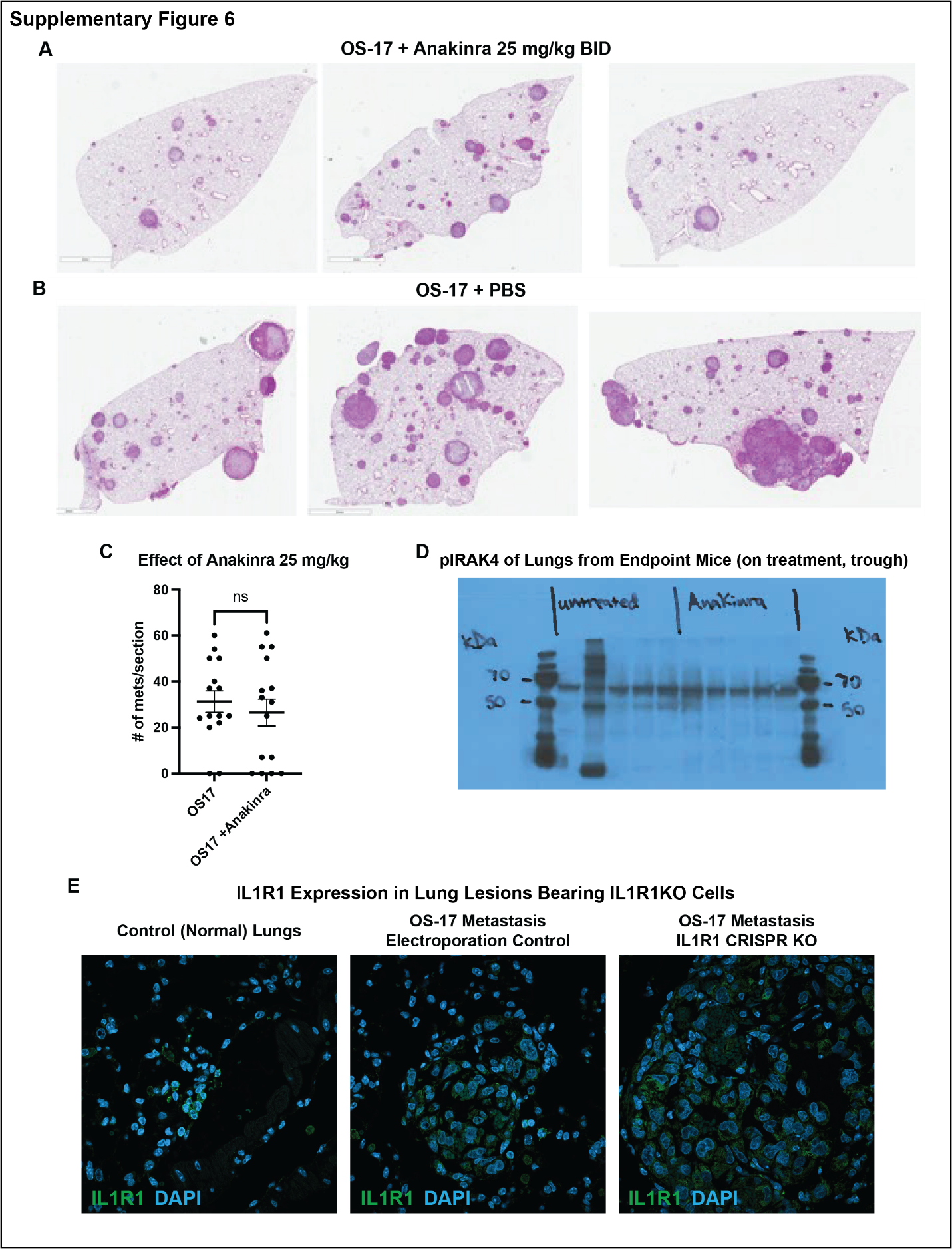

**Supplementary figure 6**. *In vivo* anakinra pharmacodynamics and IL1R1 expression in OS-17 cells *in vivo*. A-C). Mice were injected with OS-17 cells via tail vein and randomized to received PBS (vehicle) or anakinra 1 day after injection. At end point, lungs were extracted and processed for H/E to quantify metastatic lesions. No significant difference found in mice treated with 25mg/kg BID of anakinra. D) Western blot pharmacodynamics demonstrate that 25mg/kg BID did not significantly lower p-IRAK4 signal (surrogate marker for IL1R1 signaling). Immunohistochemistry of IL1R1 (stains human and mouse IL1R1) in control lung, OS-17 control metastasis, and IL1R1 CRISPR knock-out (rare) metastasis demonstrates that rare metastases in IL1R1 knock-out cell lines express IL1R1, suggesting that some IL1R1+ cells (presumably non-transduced) escaped selection.
